# Full species-wide leaf and seed ionomic diversity of *Arabidopsis thaliana*

**DOI:** 10.1101/2020.11.09.373282

**Authors:** Ana Carolina A. L. Campos, William F.A. van Dijk, Priya Ramakrishna, Tom Giles, Pamela Korte, Alex Douglas, Pete Smith, David E. Salt

## Abstract

- Soil is a heterogenous reservoir of essential elements needed for plant growth and development. Plants have evolved mechanisms to balance their nutritional needs based on availability of nutrients. This has led to genetically-based variation in the elemental composition ‘ionome’, of plants, both within and between species.
- We explore this natural variation using a panel of wild-collected, geographically widespread *Arabidopsis thaliana* accessions from the 1001 Genomes Project including over 1,135 accessions, and the 19 parental accessions of the Multi-parent Advanced Generation Inter-Cross (MAGIC) panel, all with full-genome sequences available.
- We present an experimental design pipeline for high-throughput ionomic screenings and analyses with improved normalisation procedures to account for errors and variability in conditions often encountered in large-scale, high-throughput data collection. We report quantification of the complete leaf and seed ionome of the entire collection using this pipeline and a digital tool-IonExplorer to interact with the dataset.
- We describe the pattern of natural ionomic variation across the *A. thaliana* species and identify several accessions with extreme ionomic profiles. It forms a valuable resource for exploratory QTL, GWA studies to identify genes underlying natural variation in leaf and seed ionome and genetic adaptation of plants to soil conditions.

## Introduction

The ionome represents the sum of all mineral nutrient and trace elements of biological or environmental importance in an organism (Lahner *et al*., 2003). The coupling of the analyses of the ionome with genome-enabled genetics is termed ionomics (Salt, Baxter and Lahner, 2008). Ionomics have been successfully applied in other organisms such as yeast (Eide *et al*., 2005; Yu *et al*., 2011, 2012), milk (Hadsell *et al*., 2018), human serum (Konz *et al*., 2017), mammalian organs (Ma *et al*., 2015) and human cell lines (Malinouski *et al*., 2014). Plants being sessile organisms, have evolved intricate regulatory mechanisms to balance uptake and distribution of mineral nutrients and trace elements in response to physical and chemical variation of the soil. *Arabidopsis thaliana* is a non-crop, genetic model plant species extensively used in plant biology due to its small genome size and extensive genetic resources. Ionomic studies in this species have led to the identification of genes involved in numerous processes, such as Casparian strip formation (Hosmani *et al*., 2013; Kamiya *et al*., 2015), sphingolipid biosynthesis (Chao *et al*., 2011), phloem transport (Tian *et al*., 2010), vesicle trafficking (Gao *et al*., 2017), one-carbon metabolism (Huang *et al*., 2016), and iron homeostasis (Hindt *et al*., 2017). Further, there is increasing evidence for organ-specific ionomic data to provide insights into understanding underlying functions (McDowell *et al*., 2013; Baxter *et al*., 2014; Pauli *et al*., 2018). These make ionomics on plants particularly interesting as it can reflect variation in soil mineral composition, altered function of ion transporters, the genes that control the concentration of essential or toxic elements, and processes that indirectly impact their transport (Huang and Salt, 2016).

One of the first plant genome-wide association studies (GWAS) in *A. thaliana* explored the allelic variation in a selection of 96 wild collected accessions and investigated 107 different traits, including the leaf ionome (Atwell *et al*., 2010). The study identified the gene *HKT1;1* as being associated with phenotypic variation in leaf Na concentration, validating what had previously been observed in a bi-parental mapping population (Rus *et al*., 2006). An extension of this approach using the *A. thaliana* HapMap population of 349 accessions showed improved detection and identified three causal genes underlying variation in the concentration of elements in leaves - *MOT1* for Mo, *HMA3* for Cd, *HAC1* for As (Baxter *et al*., 2008, 2010; Chao *et al*., 2012; Chao, Chen, *et al*., 2014; Forsberg *et al*., 2015) and with re-identification of *HKT1;1*. In addition, the intraspecific ionomic variation of accessions has been used with success in crops like rice *Oryza* spp. (Lou *et al*., 2015; Ricachenevsky *et al*., 2018; Yang *et al*., 2018), soybean (Ziegler *et al*., 2018) and *Brassica rapa* (Bus *et al*., 2014; Thomas *et al*., 2016). It provides a promising basis to uncover the genetic and physiological mechanisms that regulate the elemental concentrations in plants both between (Watanabe *et al*., 2007), and within species (Huang and Salt, 2016; Neugebauer *et al*., 2018), and the potentially adaptive significance of this variation (Poormohammad Kiani *et al*., 2012; Busoms *et al*., 2018). Further, it highlights the power of screening large collections of accessions for the identification of rare alleles associated with phenotypic differences in ionome. The 1001 Genomes project made available detailed genome sequence information for a large diverse collection of around 1,135 wild collected *A. thaliana* accessions from Europe, Central Asia, North Africa and North America (Weigel and Mott, 2009; Cao *et al*., 2011; Alonso-Blanco *et al*., 2016). With a threefold higher number of accessions, and a genetic diversity of more than 10 million bi-allelic SNPs, we decided to experimentally characterize the full natural ionomic variation of this population to help uncover the genetic architecture underlying the natural ionomic variation within *A. thaliana* in both leaves and seeds. The *A. thaliana* seed ionome would be a unique dataset to verify if the genetic basis of the seed ionome in *A. thaliana* and important crops such as rice, maize, soybean, etc. are the same. Further, it forms a unique dataset to explore and better understand the relatedness between the leaf and seed ionome.

Due to the size of the 1001 collection, large-scale ionomic experimental workflows were developed where plants were grown and analysed on a continuous basis over the course of several months. Previous ionomic studies on collections of natural accessions faced the challenge of suboptimal normalisation and experimental design that may have contributed to underpowered results (Atwell *et al*., 2010; Baxter, 2010; Baxter *et al*., 2012; Chao *et al*., 2012; Chao, Baraniecka, *et al*., 2014). This highlights the importance of a robust experimental design and sound normalisation of stochastic variation in experimental conditions. Normalisation methods can be divided into-control, and sample based methods (Birmingham *et al*., 2009). In most high-throughput ionomic experiments with *A. thaliana*, control based normalisation techniques have been used to filter out differences between experimental blocks (plant growth trays) (Baxter, 2010; Baxter *et al*., 2012; Chao *et al*., 2012; Chao, Chen, *et al*., 2014). Methodologically, high-throughput ionomic experiments on plants resemble high-throughput microplate assessments. For these kinds of experiments, a plethora of normalisation methods have been developed and reviewed to compensate for introduced noise into these kind of experiments (Malo *et al*., 2006; Wiles *et al*., 2008; Baryshnikova *et al*., 2010; Dragiev, Nadon and Makarenkov, 2011; Yu *et al*., 2011; Caraus *et al*., 2015; Murie *et al*., 2015). Spatial bias can be compensated for by statistical techniques such as spatial smoothing in which spatial bias is modelled based on samples and compensated for with generalised additive models (Hastie and Tibshirani, 1986) or LOESS curves (Baryshnikova *et al*., 2010; Murie *et al*., 2015). In this study, we propose an experimental design and two-step normalisation procedure for high-throughput ionomic studies, that extend previously described normalisation procedures such as REML normalisation (Broadley *et al*., 2010), by using a sample based approach to normalise for spatial variation between and within experimental blocks, and a control based approach to evaluate the normalisation.

With this work, we aimed to provide a unique referential dataset of this large natural population that forms a valuable phenotypic resource to enable large-scale GWAS to identify the genetic basis of ionomic traits. The study also explores the relationship between the leaf and seed ionome and identifies new extreme accessions that can be used to generate bi- and multi-parental populations for QTL analyses. Finally, the wide geographic range of this population, along with dense sampling at specific locations, can be a starting point for new local adaptation studies.

## Results

### Validation of experimental design

We developed and optimised a high-throughput analysis pipeline for elemental profiling of twenty-two elements - Li, B, Na, Mg, P, S, K, Ca, Cr, Mn, Fe, Co, Ni, Cu, Zn, As, Se, Rb, Sr, Mo, Cd and Pb, in leaves and seeds and present data for 1,135 accessions of *A. thaliana* from the 1001 Genomes project which includes 180 Swedish accessions and the 19 parental lines of the MAGIC population (Kover *et al*., 2009; Weigel and Mott, 2009; Long *et al*., 2013; Alonso-Blanco *et al*., 2016).

We included four normalisation lines Col-0, Cvi-0, Fab-2 and Ts-1 in our experimental design to correct for experimental variation in the ionomic profile between plant growth trays. We confirmed that the ionomic profiles of these selected accessions are distinct (**Figure S1a**). We observed that Cvi-0 showed an extreme phenotype for several elements - high Mo, As and Cd, low Sr, Ca and Mg and was the most distinct accession among the four selected. To account for variation between and within trays we included four replicates per normalisation line which were distributed evenly in growth trays on a row and column basis (**Figure S1b,c**). To account for the impact of instrumental noise caused by drift in the sensitivity of the ICP-MS on the analysis, we applied a linear model that incorporates the effect of signal drift within a given run as a covariate to compensate for variation between ICP-MS runs. Our results indicate significant drift of the analytical signal within the ICP-MS runs for all elements except Na and Zn, and the drift patterns varied between ICP-MS runs. There were also significant differences in the sensitivity of the instrument across all elements. Once the analytical signal for each element was corrected for drift, we accounted for LOD for each element on the ICP-MS by excluding quantifications of a given element in a sample that had values below 3 standard deviations of the mean value of the blanks (**Table 1**). This resulted in the exclusion of a large number of values for elements such as B (81.6%), Cr (70.9%), Ni (66.5%) and Pb (95.6%), which were therefore considered unreliable and excluded from further analyses.

**Table 1.**
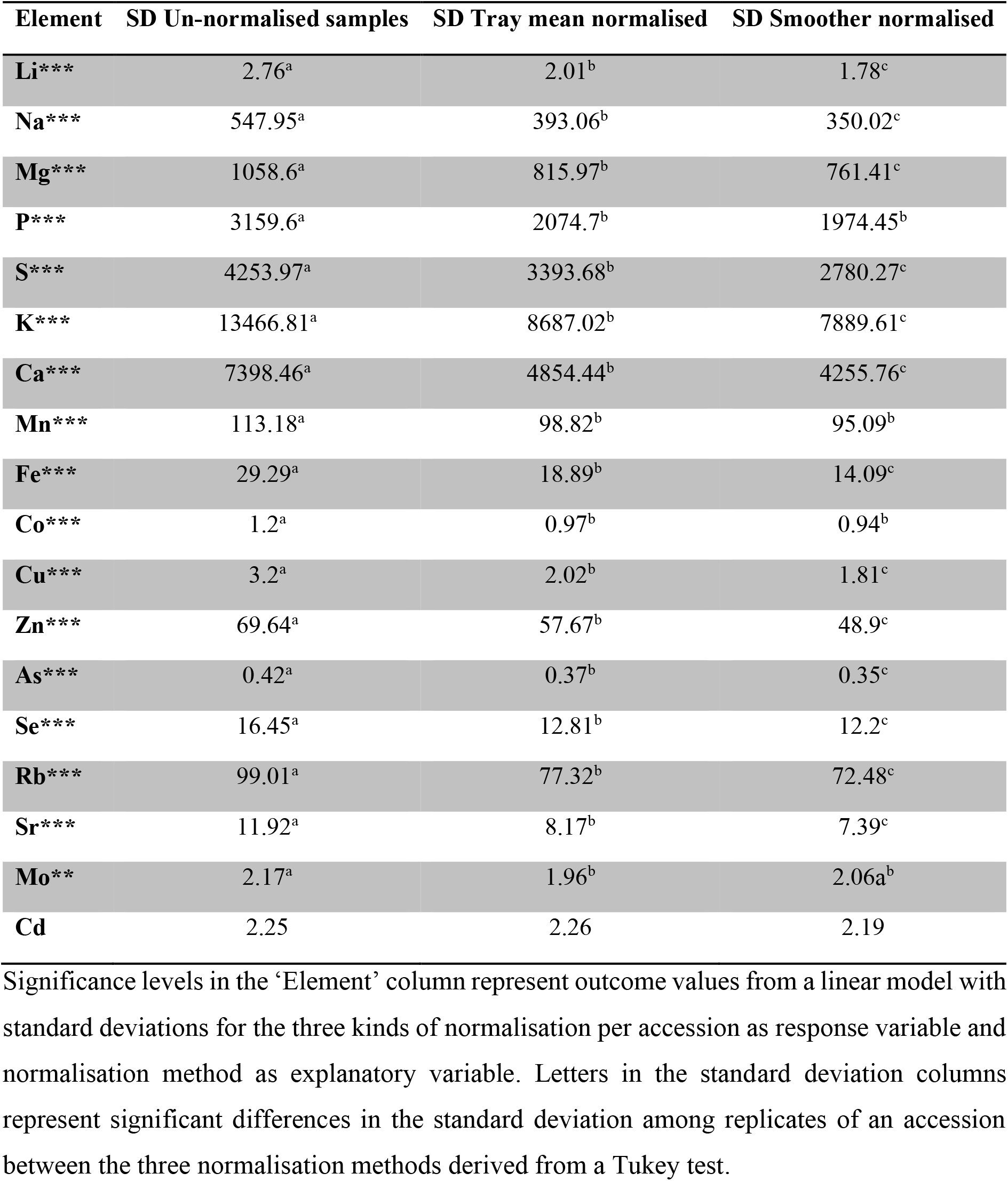
Output of the linear model comparing standard deviations from the three different normalisation approaches per accession over all the accessions studied.

### Normalisation strategies employed and validated across the pipeline

To increase the robustness of the ionomic dataset, we utilised four common check-up lines Col-0, Cvi-0, Fab-2 and Ts-1, rather than just Col-0. This provided a better fit for the data with a goodness of fit for the calculated versus measured weights of 92.3% for leaf samples and 83.6% for the seed samples. We found significant differences in elemental concentrations for all elements between the different trays used to grow the plants, as was reflected in similar patterns of variation between trays of the check-up lines (**Figure S2, S3, Table S1, S2**). The application of a tray-specific smoother using a GAM to account for the systematic noise successfully compensated for the differences within trays for all elements with no further within tray differences for either leaf or seed ionomes, except for Zn for the seed ionome (**Table S1, S2**).

### Distribution of elemental data and identification of new accessions with extreme ionome profiles

We examined the leaf and seed elemental distribution across the accessions in the study. We found high levels of variation in the leaf elemental concentrations across the accessions, which included some new, and already described extreme accessions for most elements appearing at the tail of the distribution (**Figure 1a**). We were also able to detect several accessions with extreme leaf ionome phenotypes never described before by selecting the 10 top and bottom accessions for each leaf element concentration (**Table S3**). Among the 10 accessions with highest and lowest concentrations of each element some occurred multiple times indicating a multi-element extreme ionomic profile. For some elements, these were chemical analogues, such as S and Se, K and Rb, Ca and Sr. However, even when not considering the chemical analogues, there were 10 accessions that stood out with more than three elements among these extremes namely, Stilo-1, Cvi-0, Pva-1, CS75436, Ham 1, Ala-0, Castelfed-2-201, UKID13, Ha-SB and Reg-0 (**Figure 2a**). The check-up line Cvi-0 and Fab-2 and the accession *Ler-0* from the MAGIC parents appear in this list. High variation across accessions were observed for most elements quantified in the seeds, with new and already described extreme accessions appearing in the tails of the distribution (**Figure 1b**). Several accessions with extreme seed ionome phenotypes were identified by selecting the ten top and bottom accessions for seed elemental concentration, with several being described for the first time (**Table S4**). Not considering chemical analogues, 9 accessions stood out for having more than 3 elements among these extremes-Lip-0, Pa-1, Bar-1, Aln-30, Lam-0, Urd-1, Iso-4, Ven-0 and Bela-3 (**Figure 2b**). There were 41 accessions that had 2 elements among these extremes for the leaf and seed data (**Table S3, S4**). The MAGIC parents Hi-0, Tsu-0 and Zu-0 appear in this list. We then, calculated the broad sense heritability to estimate to what extent the variation in the leaf and seed ionome across the accessions were due to genetic factors (**Table 2**). We see the heritability values being lower than reported in previous studies for the leaf ionome (Atwell *et al*., 2010; Baxter *et al*., 2012). The broad sense heritability of elemental concentration in leaves across accessions ranged from 4.32% for Zn to 77.14% for Mo. For the seed ionome, heritability values ranged from 2.07% for Ni to 36.31% for Se (**Table 2**). These findings provide evidence of the power of using larger populations, such as the 1001 genome project collection, to capture a more complete picture of the existing natural variation for traits such as the leaf and seed ionome in *A. thaliana* and its potential to be replicated in other plant species.

**Figure 1.**
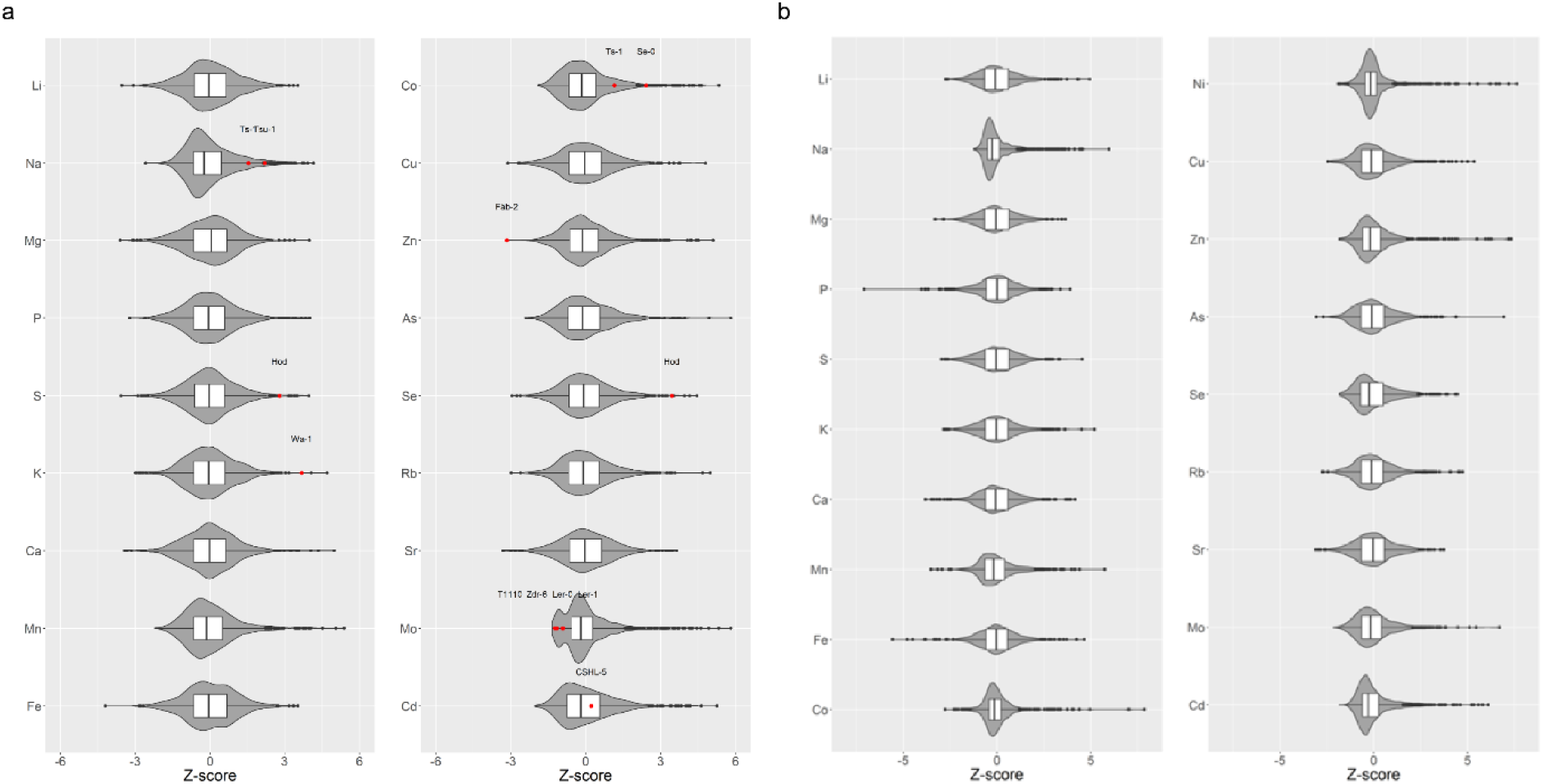
Violin plots of Z-scores indicating the distribution of the elemental levels in (a) leaves and (b) seeds across the analysed *Arabidopsis thaliana* accessions. Highlighted in red are accessions from previous studies with extreme values for a particular element concentration.

**Figure 2.**
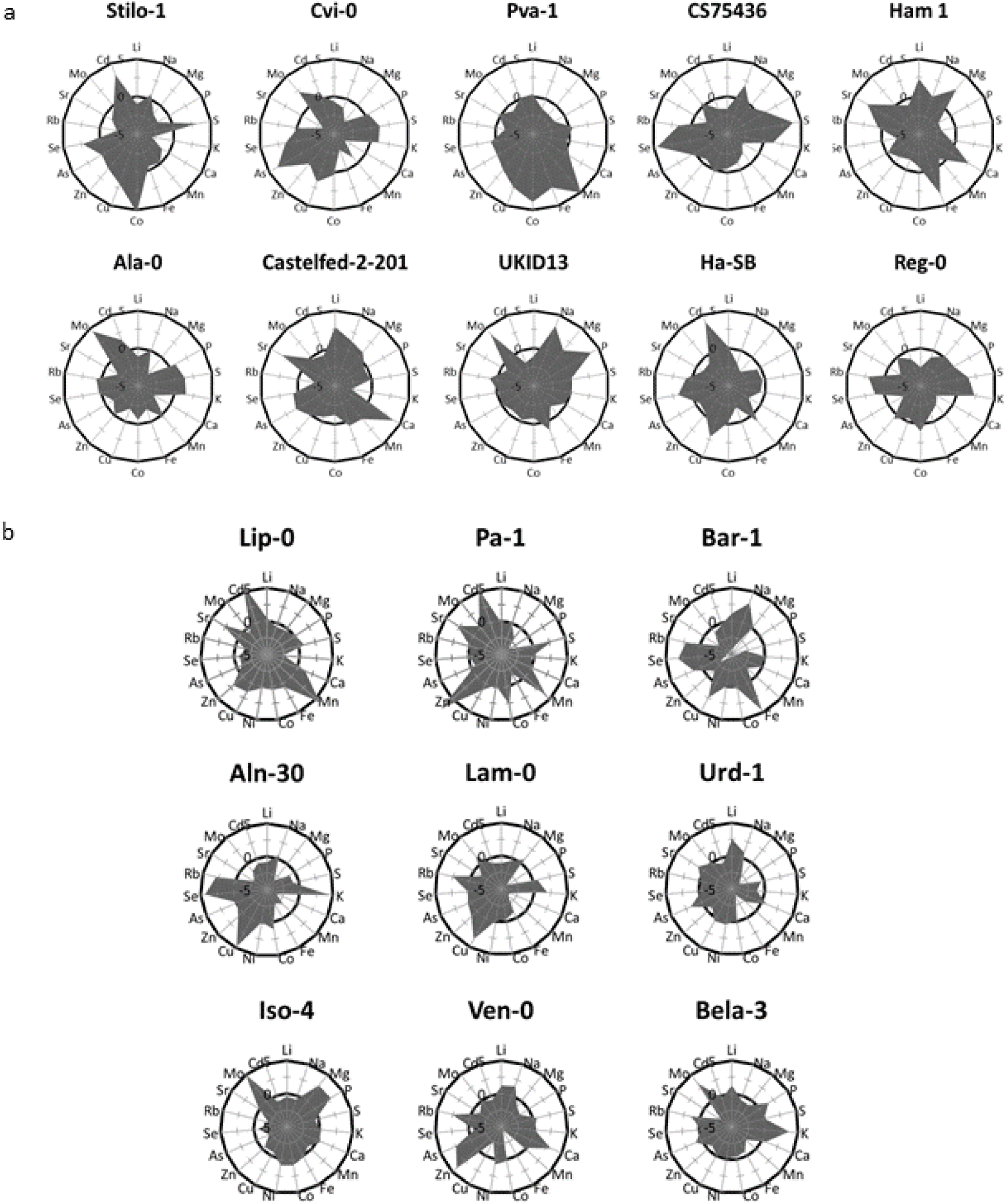
Radar plots with the Z-values of the studied elements in the (a) ten leaf multi-element and (b) nine seed multi-element extreme accessions. Axis display Z-scores calculated per element.

**Table 2.** Summary statistics of elements concentration in the leaves and seeds of the *A. thaliana* accessions used in the study.

|  | Leaf ionome |  |  |  | Seed ionome |  |  |  |
| --- | --- | --- | --- | --- | --- | --- | --- | --- |
| **Element** | **Mean** | **SD** | **RSD** | **Heritability** | **Mean** | **SD** | **RSD** | **Heritability** |
| **Li** | 14.48 | 0.63 | 4.38 | 16.58 | 2.13 | 0.54 | 25.42 | 29.77 |
| **Na** | 1163.82 | 391.41 | 33.63 | 52.89 | 45.50 | 30.37 | 66.74 | 18.36 |
| **Mg** | 4819.69 | 402.21 | 8.35 | 26.49 | 1419.39 | 96.85 | 6.82 | 26.09 |
| **P** | 16092.66 | 1210.18 | 7.52 | 28.67 | 5185.68 | 179.92 | 3.47 | 11.33 |
| **S** | 12820.45 | 1231.15 | 9.60 | 21.17 | 3652.94 | 375.00 | 10.27 | 9.39 |
| **K** | 71770.74 | 6679.26 | 9.31 | 43.54 | 6784.59 | 664.47 | 9.79 | 26.31 |
| **Ca** | 41517.42 | 2431.91 | 5.86 | 25.47 | 4348.66 | 341.86 | 7.86 | 20.41 |
| **Mn** | 229.35 | 12.23 | 5.33 | 5.48 | 25.97 | 5.80 | 22.33 | 32.92 |
| **Fe** | 139.95 | 3.26 | 2.33 | 10.49 | 19.61 | 0.85 | 4.34 | 6.26 |
| **Co** | 2.50 | 0.60 | 23.90 | 29.38 | 0.29 | 0.04 | 13.97 | 3.94 |
| **Cu** | 7.82 | 0.63 | 8.12 | 16.68 | 2.75 | 0.23 | 8.47 | 16.45 |
| **Zn** | 92.42 | 6.48 | 7.01 | 4.32 | 21.81 | 2.89 | 13.24 | 6.42 |
| **As** | 0.99 | 0.08 | 8.32 | 9.98 | 1.68 | 0.19 | 11.06 | 17.66 |
| **Se** | 73.26 | 9.56 | 13.05 | 39.87 | 11.39 | 8.18 | 71.84 | 36.31 |
| **Rb** | 406.14 | 43.66 | 10.75 | 28.39 | 41.67 | 4.58 | 10.99 | 24.96 |
| **Sr** | 66.53 | 4.25 | 6.38 | 28.34 | 5.91 | 1.05 | 17.84 | 29.94 |
| **Mo** | 6.23 | 4.44 | 71.29 | 77.14 | 1.30 | 0.51 | 39.48 | 34.37 |
| **Cd** | 2.46 | 0.24 | 9.88 | 4.95 | 0.15 | 0.13 | 87.28 | 33.94 |

The parameters shown are: mean concentration (ppm), standard deviation (SD, ppm), relative standard deviation (RSD, %) and broad sense heritability (%).

### Analysis of the leaf and the seed ionome

#### Elemental correlation analysis

To understand the relationship between the elemental concentrations, we performed a correlation analysis and grouped them using K-means clustering (**Figure 3a**). For the leaf ionome, the elements grouped into ten clusters with a stronger association among elements within a cluster than between clusters. As expected, the chemical analogues S and Se, K and Rb, Ca and Sr, had the highest correlation and grouped in the same clusters. The elements Li, Fe, Ca and Sr clustered together and were positively correlated between themselves, but negatively correlated with K, Rb, S and Se. Although not in the same cluster, some of these elements also showed a positive correlation with Mg. Other clusters of elements positively correlated were Cu, Zn, Mn and Cd. The elements Mg, Co, Mo and Na did not cluster with any other elements, and Mg alone showed moderate correlation with other elements. For the seed ionome, elements grouped into nine clusters and the only pair of chemical analogues we found in the same cluster with strong positive correlation were K and Rb (**Figure 3b**). S and Se, and Ca and Sr grouped in separate clusters, but positively correlated. The first large cluster was composed of K, Rb, Cu and Se, all positively correlated with each other. Most of these elements also showed a negative correlation with Sr, Mn, Ca and Fe. In the second largest cluster we find the positively correlated elements Zn, Sr, Mn and Cd and most showed moderate negative correlation with Mg. Other clusters of elements identified showed weak to moderate correlations with each other.

**Figure 3.**
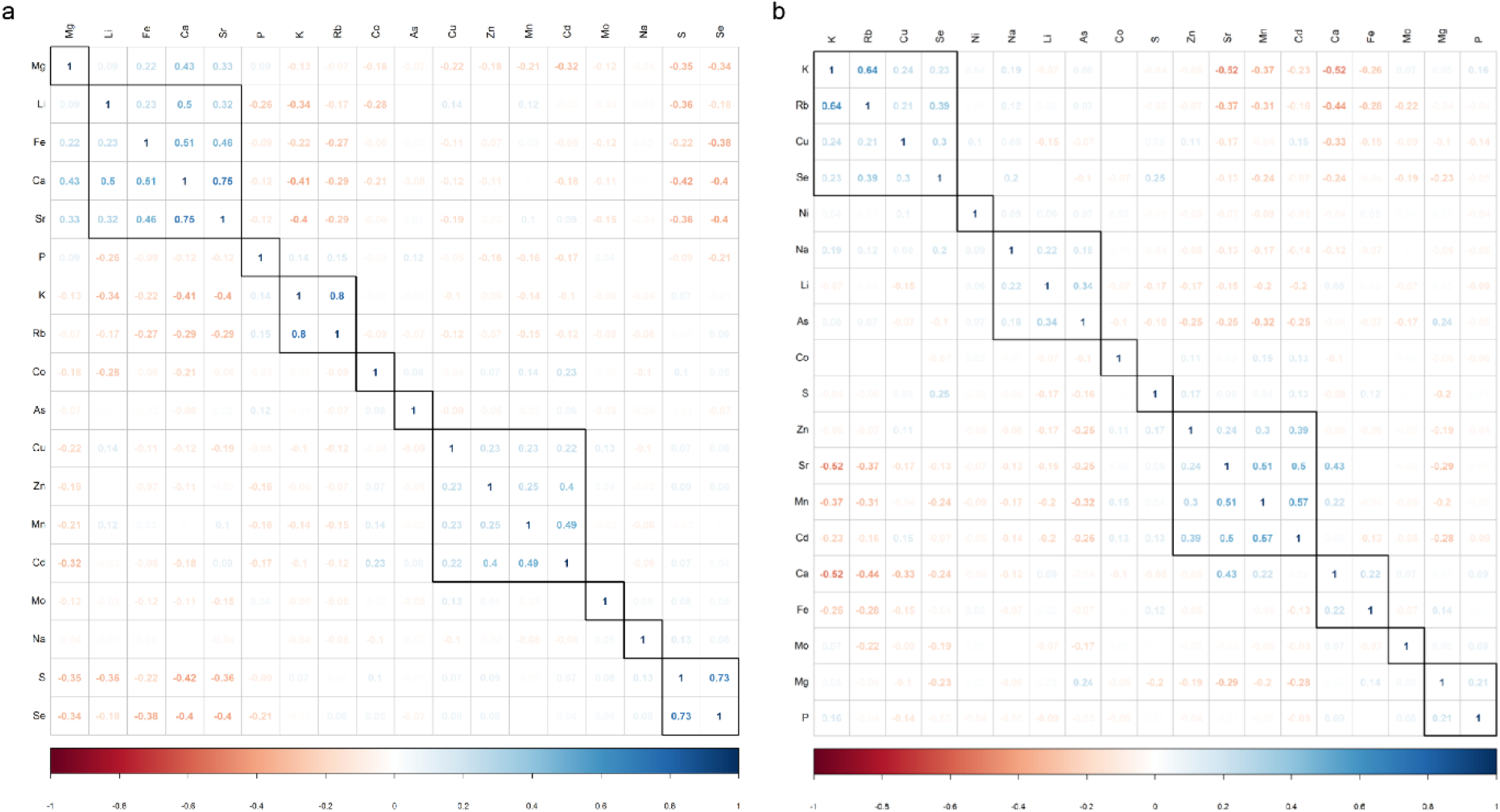
Correlogram with elements concentrations across the studied *A. thaliana* accessions in the leaves (a) and seeds (b). Blocks indicate clusters of elements based on unassisted hierarchical clustering. Number of clusters are defined based on K-means clustering.

#### Principal Component Analysis

Given that different soils and environments may drive differences in the ionome, one might expect the ionome of accessions to reflect their geographical origins. In order to verify the existence of these patterns, we performed a Principal Component Analysis (PCA) on the leaf and the seed ionome as it highlights extreme phenotypes taking into account the ionome as a whole. First, the accessions were grouped based on similarities and differences in the ionome. Then for signs of regional adaptation, they were then grouped based on the geographical groupings described in a previous study on these accessions (Alonso-Blanco *et al*., 2016). For the leaf ionome, the first two principal components (PCs) together explained only 34.5% of the total variation in the leaf elements across the studied accessions (**Figure S4a**). The PCA largely supports the clustering of the different elements found in the elemental correlation analysis. The chemical analogues S and Se, K and Rb, Ca and Sr, and elements Cd, Zn and Cu are grouped together, likely due to low specificity of some uptake and transport proteins for these elements. The first PC explains 21% of the variation, mainly driven by variation in the concentration of Cd, Mn, Zn, Cu and P. The second PC explains 13.5% of the variation in the leaf ionome, largely determined by the chemical analogues Ca and Sr, Se and S, K and Rb, and Fe, Mg and Li. Some elements had opposing loadings in the PCA plot which could be indicative of a crosstalk between their homeostasis. The notable ones were P vs Mn and Mg vs Co. However, no strong correlation was found between these pairs of elements (**Figure 3a**). Five of the ten multi-element extreme accessions identified stand out as extremes in the PCA plot (**Figure S4a**). However, the PCA was unable to identify strong signs of regional identity based on the geographical groups. Some extreme accessions for the same element that stand-out in the PCA, were from the same geographical group: Men-2 and Vas-0 from Spain with high Se; Panik-1 and Borsk-2 from Asia with high Sr and Ca, respectively. Interestingly, although the accessions Castelfed come from the same region and sampled very close to each other, they do not have a similar leaf ionomic profile, and do not cluster together in the PCA. For the seed ionome, we found that the first two PCs together explained 31.3% of the total variation in the seed ionome across the accessions (**Figure S4b**). The first PC explained 18.4% of the variation, mainly driven by variation in the concentrations of Ca, Mg, Cu and Cd. The second PC explained 12.9% of the variation, largely determined by the elements Sr, Mn and the chemical analogues K and Rb. Indications of crosstalk were observed for Fe and Mo *vs* Cu and Mg, Zn and Co *vs* Li and As *vs* Cd. However, only As had a clear negative correlation with Cd. The accessions Lam-0 and Bar-1, previously identified as extremes for multiple elements, likewise stand out as extremes in the seed ionome PCA.

### Clustering of accessions based on the ionome and evidence of local adaptation

To examine possible signs of regional adaptation, first we grouped the accessions based on similarities and differences in the ionome against the above-mentioned geographical groupings (Alonso-Blanco *et al*., 2016). In general, neither the leaf or seed ionome showed distinct clustering based on the geographical groups either looking at the total set of accessions or the 10 top and bottom extreme accessions (**Figure S5 a,b; Figure S6 a,b**).

#### Geographic distribution of extreme accessions per element

To further explore the possible relationship between geographic regions and the leaf and seed ionome, we looked at the geographic distribution of the 10 top and bottom extreme accessions for each leaf and seed element concentration. The geographic location of the original collection site and the leaf ionomic phenotype of these extreme accessions were visualised on a map (**Figure 4 a,b**). We could confirm two clusters of accessions from North and South Sweden with a common extreme leaf ionome phenotype that also clustered in the heatmap (**Figure S6a**). Accessions with extremely low K concentrations occurred together in the northern region of Sweden, and accessions with extremely low S concentrations occurred close to each other in the southern region of Sweden. The clusters identified with this approach consisted of accessions that occurred together in Azerbaijan at the Caspian Sea with extremely high S concentrations in leaves, at Castelfed with high Cd and Sr, and Spain with high K and Rb (**Figure 4 a,b**). While using the same approach for the seed ionome we also identified new clusters of accessions not found with the heatmaps. A cluster of accessions from Spain, with extremely high K and low Sr concentration in seeds, and several accessions with extremely high Se and low Zn concentration in seeds distributed across Spain (**Figure 4 c,d**). Accessions with extremely low Ni concentration in seeds that occurred together in the UK and accessions with extremely low Cd occurring in the Southeast of Sweden and in Azerbaijan at the Caspian Sea (**Figure 4 a,b**). Finally, we could also confirm the cluster of accessions from the South of Sweden with extremely low Se concentration in seeds found in the heatmaps (**Figures S5b and S6b)**. We conclude, based on these observations that the high-throughput ionomics approach using a largescale worldwide collection was able to identify novel geographical patterns of variation based on analysis of the leaf and seed elemental concentrations.

**Figure 4.**
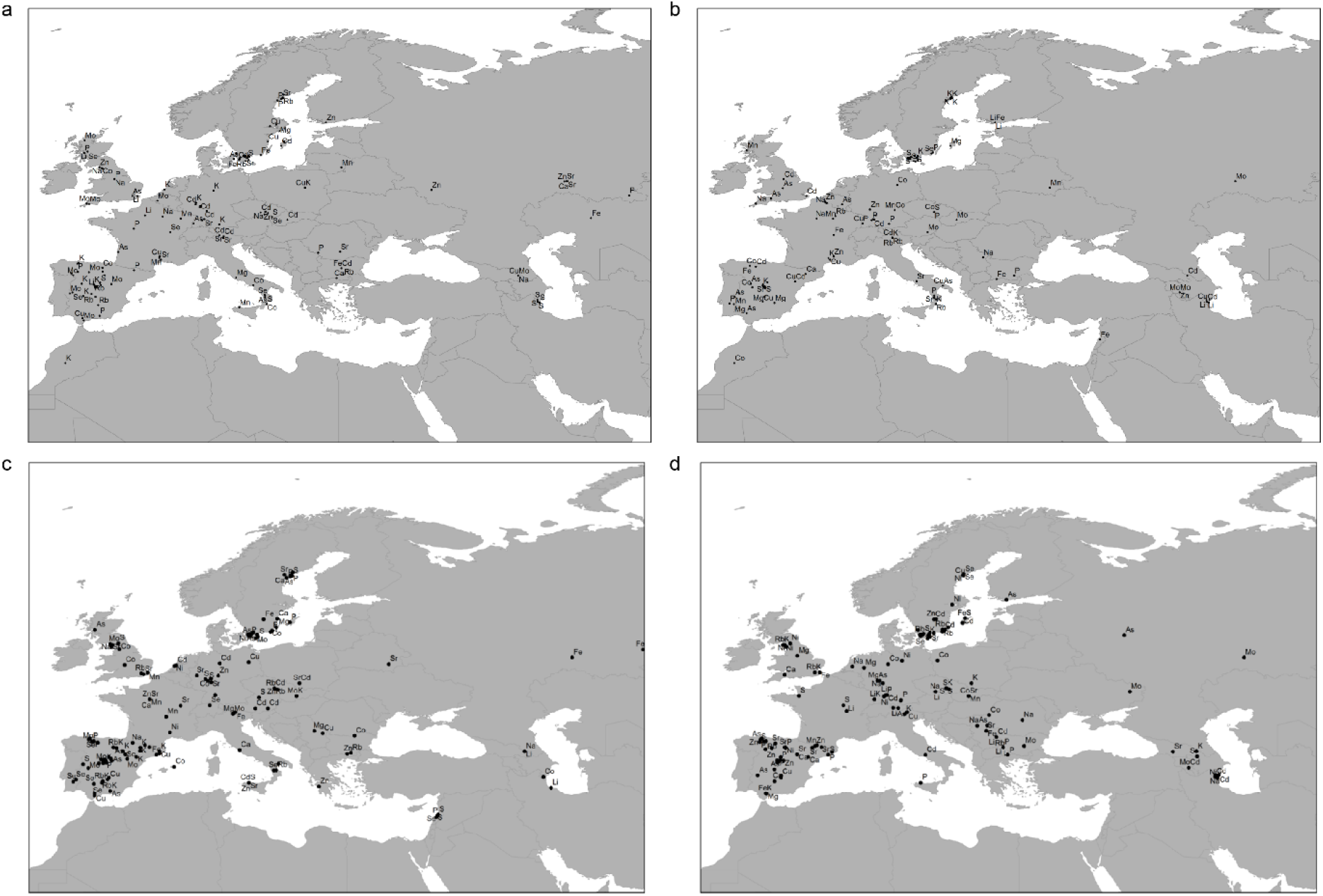
Geographic distribution of the ten *A. thaliana* accessions with the top (a) and bottom (b) leaf and 10 top (c) and bottom (d) seeds extreme phenotypes for each element analysed in this study (For overview of the accessions see **Table S3, S4**)

### Comparative analysis between different tissues - the leaf and seed ionome

#### Correlation analysis of the leaf and seed elemental profiles

We explored the relationship between the concentration of different elements in different tissues of the plant across all accessions. The elements grouped into 13 clusters with a stronger relationship between elements within the same cluster (**Figure 5**). Clusters identified when considering the seed and leaf ionomic data together largely support the clusters also identified when analysing the leaf and seed tissues independently. The exceptions however were S, Mo and Na whose seed and leaf concentrations are in the same cluster and are positively correlated.

**Figure 5.**
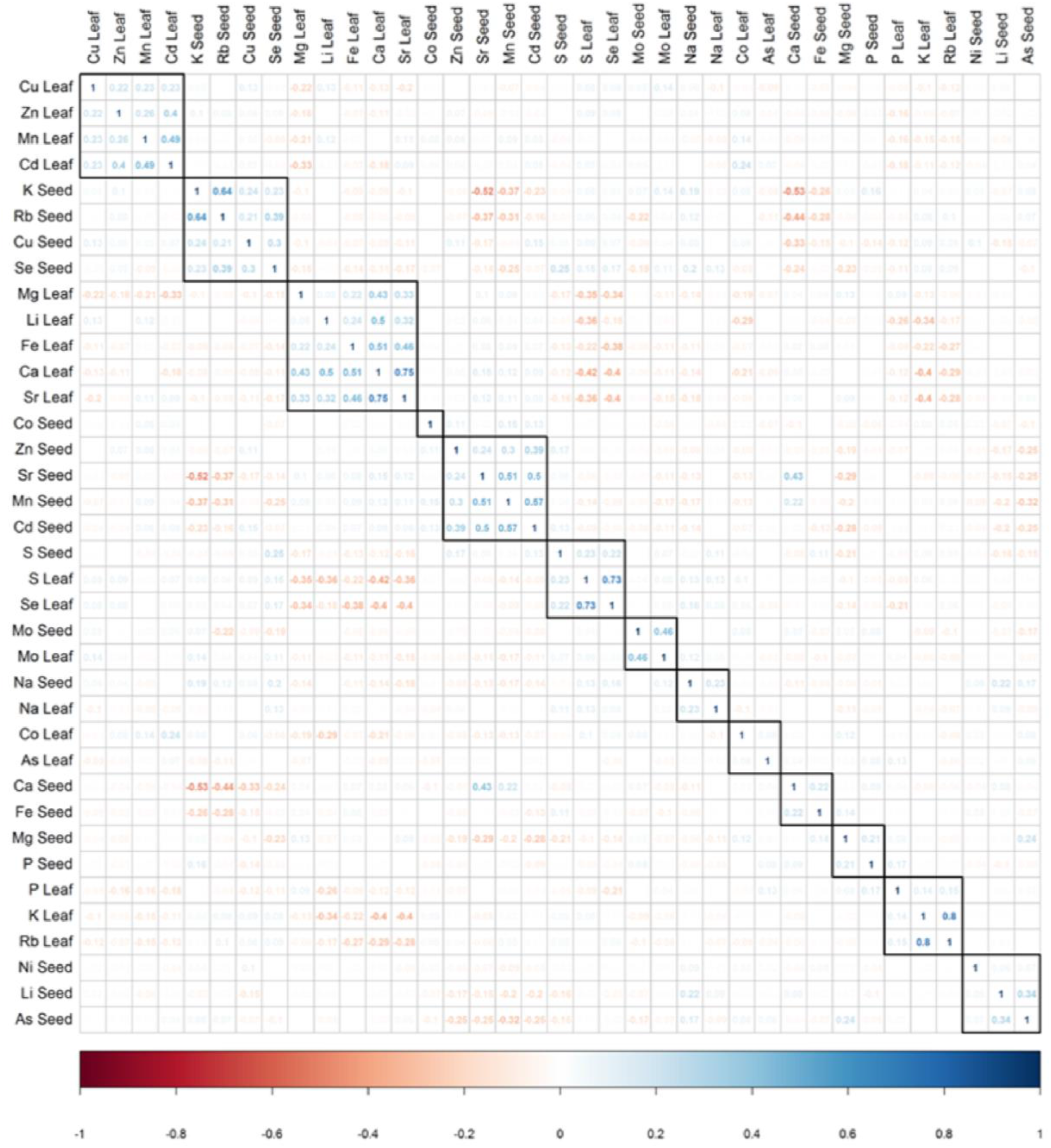
Correlogram with elements concentrations in leaves and seeds across the studied *A. thaliana* accessions. Blocks indicate clusters of elements based on unassisted hierarchical clustering. Number of clusters are defined based on K-means clustering.

#### PCA analysis of the leaf and seed ionome combined

To examine the relationship between the leaf and seed ionome, we performed a PCA analysis on the element concentrations of the two tissues. The first two PCs together explained 20.6% of the total variation observed and give an indication of accessions with extreme ionomic phenotypes for both tissues besides supporting some of the clusters of the different elements previously described in this paper (**Figure 6**). The first PC explains 12% of the variation and is mainly driven by Mn, Cd, Sr and K concentrations in seeds, while the second PC explains 8.6% of the variation largely driven by S, Se, Ca and Sr concentrations in leaves. Even though the PCA analyses takes the ionome for both tissues as a whole, elements with opposing loadings, which indicate possible crosstalk for homeostasis, were only found for the same tissue. Several elemental concentrations in both seeds and leaves had extremely low PCA loadings. Most of these were the ones that correlated weakly with other elements, both when considering the seed and leaf ionome together and independently.

**Figure 6.**
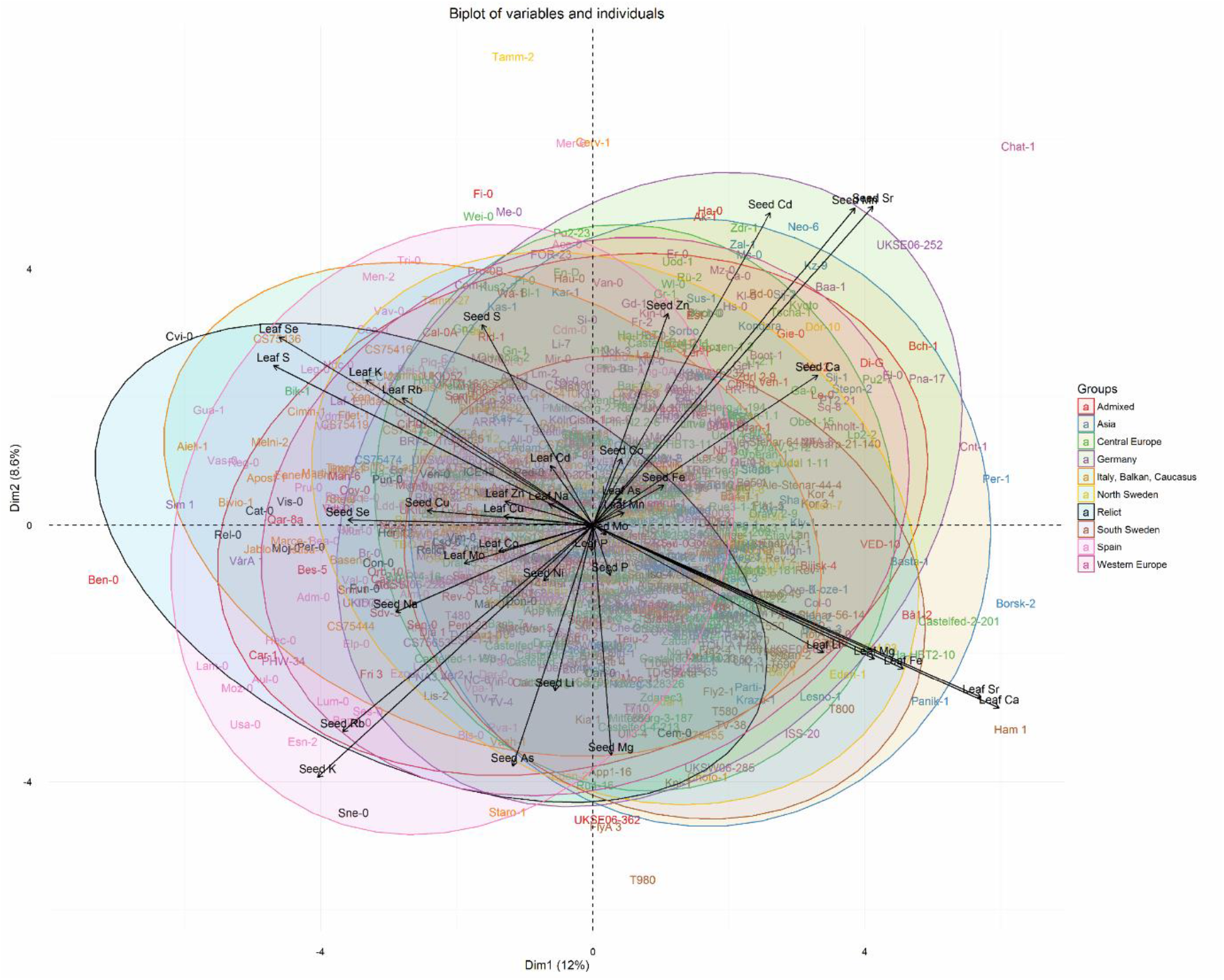
Biplot of the PCA analysis of both the leaf and seed ionome of all the studied *A. thaliana* accessions with the regional grouping as described in (Alonso-Blanco *et al*., 2016).

#### Identification of accessions with extreme profiles in both leaf and seed ionome

Although not all elements in the ionome showed a strong relationship between the leaf and the seed ionome, we were able to identify several accessions with extreme leaf and seed ionome profiles. The extreme accessions identified were separated in two groups: (i) Accessions with an extreme phenotype for the same element or chemical analogues in both seed and leaf (**Figure 7a**), and (ii) Accessions which are among the ten top and bottom accessions for each element concentration in both leaves and seeds across the accessions in the study (**Figure 7b**). Four out of these accessions also stood out as seed and leaf extremes in the PCA plot: T980; Sim1, Lam-0 and Castelfed-2-201.

**Figure 7.**
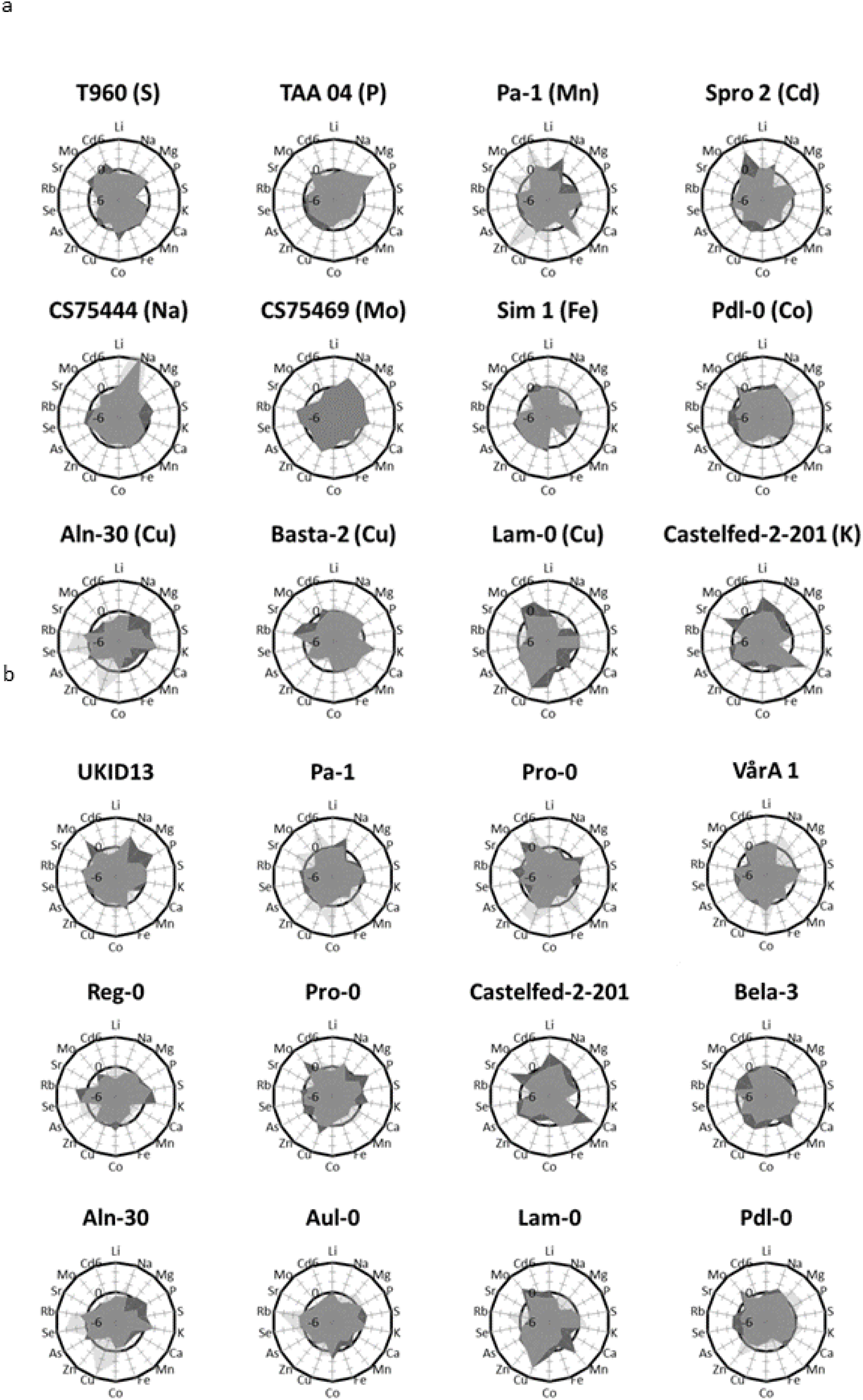
(a) Accessions with an extreme phenotype for the same element or chemical analogues in both seeds and leaves; (b) Accessions which are among the ten top and bottom for each element concentration in both leaves and seeds across all studied *A. thaliana* accessions.

### Ion Explorer – Interactive data viewer to explore the leaf and seed ionome

To facilitate convenient interaction with the complete species-wide leaf and seed ionomic data, we present an interactive web-based tool made available online: Ion Explorer https://ffionexplorer.nottingham.ac.uk/ionmap/. Ion Explorer allows interactive visualisation, analysis and comparison of the two large datasets. The Ion Map section provides a one-stop preview to the users of the range of the accessions across the globe and the elemental levels of each element in the leaf and seed. The data can be filtered for various parameters based on leaf/seed dataset(s), element(s), location, and/or accession(s) of interest. The filtered data are reflected in the map as well as transferred for basic comparative analysis to the ‘Graph’ section of the tool. At present we have provided the functionality to compare the z-scores of all elements of select accessions, correlation and PCA that considers the filtered data as a whole. One can interact with the leaf and seed data independently and in parallel. The complete and filtered dataset can be downloaded as a .csv file, while the maps, correlograms, radar and PCA plots generated can be downloaded as image files. Further, the user can also upload data to the tool to perform comparative analyses against the dataset. The user functionality of the tool is constantly being updated.

## Discussion

To facilitate the development of a fuller picture of the genetic basis underlying natural variation in the ionome we present the largest available dataset to date of the leaf and seed ionome of over 1,135 accessions of *A. thaliana* from the 1001 Genome project (Kover *et al*., 2009; Weigel and Mott, 2009; Long *et al*., 2013; Alonso-Blanco *et al*., 2016).

### Experimental design and data analyses of high-throughput ionomic profiling experiments

There have been ongoing efforts to optimise the efficiency of ionomics workflows and the quality of the phenotypic data produced. In this study, we present an improved method for large-scale ionomic studies, including experimental design and data analysis. Large-scale ionomic experimental workflows where plants are grown and analysed on a continuous basis often span several months. Given the length of time required to grow and analyse thousands of plants, even with best efforts to maintain standard conditions, variation in environmental conditions (temperature, watering, etc.) between experiments are unavoidable. Therefore, it is essential to design the experiment to include multiple replicates of each accession spread across multiple plant growth trays to take into account the interaction between tray and accessions, which is what we have done in this study. Our observations emphasise the need for randomisation and robust normalisation strategies to reduce type I (false positive) and II (false negative) errors. Spatial bias introduced during plant growth can be compensated by statistical techniques such as spatial smoothing in which spatial bias is modelled and compensated with generalized additive models (Hastie and Tibshirani, 1986), or LOESS curves (Baryshnikova *et al*., 2010; Murie *et al*., 2015). Previous studies on data normalisation have found that spatial smoothing techniques were able to properly model this type of noise (Baryshnikova *et al*., 2010; Murie *et al*., 2015). In this study, we successfully expanded the approach to plant ionomic research and were able to better compensate for experimental noise than previous high-throughput ionomic screenings applying a control based normalisation approach without modelling within tray differences both for *A. thaliana* (Atwell et al., 2010, Baxter et al., 2008, 2010; Chao et al., 2012, 2014b; Forsberg et al., 2015) as well as for other crops (Lou et al., 2015; Ricachenevsky et al., 2018; Yang et al., 2018).

### Heritable ionomic variation

In genetic studies, the broad sense heritability values indicate the relative importance of genes and environment to the phenotypic variation of traits within and across populations (Visscher, Hill and Wray, 2008). Here, the heritability of variation in the leaf ionome in general are lower than previously reported for most elements except Mo. Although lower, we found correlations (0.62-0.86) between our observed heritability values and previous studies with 96 accessions, or the 360 HapMap population (Atwell *et al*., 2010; Baxter, 2010; Chao, Chen, *et al*., 2014), and two RIL populations (Buescher *et al*., 2010). For the seed ionome, even though we observed high variability, our heritability values were in general lower than those reported in previous studies with *A. thaliana* (Baxter *et al*., 2012) and other crops (Baxter *et al*., 2014; Lou *et al*., 2015). This highlights the consistency in the general trend of elements with high and low heritability values. The possible cause for the lower heritability values in our studies could be the randomisation of replicate accessions across the growth trays implemented in our experiment design but not in previous studies (Atwell *et al*., 2010; Baxter *et al*., 2010; Buescher *et al*., 2010; Chao, Chen, *et al*., 2014).

### Extreme ionomic accessions – known and new

The use of a larger population from geographically dispersed locations was able to identify several new single- and multi-element extreme accessions not reported previously (**Table S3, S4, Figure 2, 7**). This demonstrates the ability of the dataset to capture a more complete picture of the existing natural variation of the ionome. Previously identified *A. thaliana* accessions with an extreme ionomic phenotype have been used in studies to generate bi-parental mapping populations, and identify key genes involved in the homeostasis of Cd (Chao *et al*., 2012), Co (Morrissey *et al*., 2009), Na (Rus *et al*., 2006; Baxter *et al*., 2010), Mo (Baxter *et al*., 2008), As (Chao, Chen, *et al*., 2014), S and Se (Chao, Baraniecka, *et al*., 2014), and Zn (Chen *et al*., 2018) in plants. New accessions with extreme ionomic phenotypes identified in this study form an important starting point for the development of new bi-and multi-parent experimental populations for further genetic studies. Their unique ionomic phenotypes can aid the study of multi-element nutrient transport pathways, its genetic basis, and how plants respond to different growth conditions and nutritional stresses.

### Biological insights from multi-element species-wide ionomic data

The relationship and behaviour of multiple elements in the plant can be determined by three potential processes-chemical analogues, elemental transporters, or common biological mechanisms that regulate the concentration of several elements at the same time (Vreugdenhil *et al*., 2004; Salt, Baxter and Lahner, 2008). Strong correlations and relationship were observed in the leaf ionome of K and Rb, Ca and Sr, S and Se, Li, Fe and Mg among the accessions. K-Rb, Ca-Sr, S-Se are chemical analogues, and the strong correlations can be attributed to lack of selectivity in the uptake and transport of these elements in the plant (White, 2001). On the other hand, some elements, such as Na, Mo, Co and As were weakly, or not correlated with other elements, and had extremely low PCA loadings indicative that their homeostasis is highly element specific. Leaf concentrations of Mn, Cd, Cu and Zn across the studied accessions showed strong associations between these elements which has been previously described and considered to be likely a result of shared transporter proteins. For example, transporters of the NRAMP family show affinity to the elements Mn, Zn, Cd and Cu (Oomen *et al*., 2009; Cailliatte *et al*., 2010). *IRT1*, primarily identified as a Fe transporter, can also uptake Mn, Zn, Co and Cd (Korshunova *et al*., 1999; Connolly, Fett and Guerinot, 2002); *ZIP4* for Zn and Cu (Wintz *et al*., 2003); *HMA3* for Cd, Zn, and Co (Hussain *et al*., no date; Morel *et al*., 2008; Chao *et al*., 2012) and *HMA4* for Cd and Zn (Hussain *et al*., no date; Mills *et al*., 2003, 2005; Verret *et al*., 2004). The seed ionome dataset could form the basis to uncover mechanisms regulating mineral nutrients and trace elements concentration in the seed. The lack of knowledge in this field is one of the major constraints to the development of biofortified grains with high concentration of essential mineral nutrients such as Fe, Zn, Ca and Se, and avoid the sometimes consequent, accumulation of trace elements such as As and Cd (Mendoza-Cózatl *et al*., 2011; Waters and Sankaran, 2011; McDowell *et al*., 2013; Yoneyama, Ishikawa and Fujimaki, 2015; Bouis and Saltzman, 2017). Although, within the same tissue several elements were correlated, across the leaf and seed only S, Mo and Na were positively correlated, consistent with observations from a previous study using 96 accessions (Baxter *et al*., 2012). This suggests that although the seeds depend on vegetative tissue as a source of nutrients, the regulatory controls over the seed and leaf ionome are distinct.

### Evidence of species-wide local adaptation

Although several genes have been identified to regulate the ionome in *A. thaliana*, far less is known about their role in local adaptation (Huang and Salt, 2016). Some of the detailed studies on the influence of soil or climate on the *A. thaliana* ionome have been related to saline soils where they show the ability of these plants to accumulate Na in leaves based on their distance from the coast/salinity level (Rus *et al*., 2006; Baxter *et al*., 2010; Busoms *et al*., 2015), or to explain Mo homeostasis based on Mo levels in native soils of West Asian accessions (Poormohammad Kiani *et al*., 2012). Our attempts to identify associations between the leaf and seed ionome and geographic regions did not lead to any clear trends. Previous local adaptation studies have highlighted the heterogeneous nature of soils and have suggested local adaptation is likely to be found at a small to very small geographic scale (Macel *et al*., 2007; Busoms *et al*., 2015). Also the ecology of *A. thaliana* as a ruderal species often occurring in urban environments with a large heterogeneity in local soil composition may contribute to that (Guilland *et al*., 2018; Hovick and Whitney, 2019; Takou *et al*., 2019). The most prominent signals were found in the south of Sweden and at the Caspian Sea shore with accessions from these areas having extreme low leaf S and Se. The presence of a large regional collection of 180 accessions from Sweden classified into regional groups as north and south Sweden, may have contributed to our observation of an apparent association between location of origin and the leaf ionome of some Swedish accessions. New accessions identified in this study with not only high, but also low elemental concentrations in leaves and seeds hold potential for further study to identify novel alleles or genes involved in element homeostasis and local adaptation.

Taken together, this study presents a robust experimental design pipeline that allows the study of 1000’s of accessions with a normalisation strategy that deals with systematic noise within and between plant growth trays and accessions. The Ion Explorer tool holds potential to become a comprehensive resource for visualisation and interpretation of ionomic data. We envisage addition of further datasets and views to perform comparative analysis with other experimental conditions and links to other databases to increase functionality and output. The power of the presented ionomic dataset lies in the ability to undertake large-scale genetic studies, including GWAS and linkage mapping using recombinant populations, to identify new genes and alleles that regulate the plant ionome. The use of the newly identified extreme elemental accessions to generate bi- and multi-parent populations will help pave the way to uncover nutrient transport pathways in the plant and its response to different growth conditions and nutritional stresses. Finally, we present indications that local adaptation of plants to soil conditions are reflected to a limited degree in their ionome, mostly at small geographical scales. The identification of promising regions would be an interesting focus for future local adaptation studies. Such information can help improve our understanding of how plants efficiently acquire mineral nutrients from the soil and adapt to their local soil habitat.

## Materials and Methods

### Plant materials

The plants used for leaf and seed ionomic analysis included 1,135 accessions of *A. thaliana* from the 1001 Genomes project which also includes 180 Swedish accessions and the 19 parental lines of the MAGIC population (Kover *et al*., 2009; Weigel and Mott, 2009; Long *et al*., 2013; Alonso-Blanco *et al*., 2016). Seeds were kindly provided by Magnus Nordborg at the Gregor Mendel Institute of Molecular Plant Biology.

### Experimental design and growth conditions

#### Leaf ionome

Plants were grown in 32mm 7C Jiffy-7^®^ peat pellets (http://www.jiffypot.com/) pre-soaked in a solution (104 pellets in 2.6l) containing 0.142 μM Na2HAsO4•7H2O, 5.935 μM Cd(NO3)2•4H2O, 0.534 μM Co(NO3)2•6H2O, 0.053 μM LiNO3, 0.534 μM Ni(NO3)2•6H2O, 0.427 μM K2SeO4, 0.106 μM Sr(NO3)2 (98%), 0.0774 μM RbNO3 to provide these elements in trace amounts to enable detection in the plants. The final concentration of these elements in dry soil were 5.6 ppm As, 0.2 ppm Cd, 1.1 ppm Co, 1.3 ppm Li, 1.1 ppm Ni, 1.9 ppm Se, 8.7 ppm Sr and 11.7 ppm Rb. The Jiffy-7^®^ peat pellets were evenly distributed in trays with 104 places, and the trays lined with capillary matting soaked in 1.5L of 18 MΩ deionized water. Three seeds of each *A. thaliana* accession were sown in the pre-soaked Jiffy-7^®^ pellets and stratified at 4°C in the dark for 1 week to help synchronize germination. Germinated seedlings were thinned to one after one week. A growth pipeline was established with one tray sown per day, and after four weeks, one tray harvested per day. Each tray had 104 plants distributed randomly of which 88 were a single replicate of different accessions and four normalisation lines replicated four times each (**Figure S1b**). The accessions used for normalisation lines were Col-0, Cvi-0, Fab-2 and Ts-1. Six replicates of each accession were grown, each in a separate tray comprising 120 plant growth trays grown over a 37 week period. Plants were cultivated in a controlled growth room with 10 h days (~100 μmol photons m^-2^s^-1^) /14 h night, and a temperature of ~ 25°C day and 17°C night. The trays were bottom-watered 3 times in total on days 10, 17 and 24 with 0.25X Hoagland solution (1.5 mM KNO_3_, 1 mM Ca(NO_3_)2•4H_2_O, 0.5 mM MgSO_4_•7H_2_O, 0.25 mM NH_4_H_2_PO_4_, 11.5 μM H_2_BO_3_, 1.3 μM MnCl_2_•4H_2_O, 0.2 μM ZnSO_4_•7H_2_O, 0.075 μM CuSO_4_•5H_2_O, 0.028 μM MoO_3_) and Fe supplied as 10μM Fe-HBED N,N’-di(2-hydroxybenzyl)ethylenediamine-N,N’-diacetic acid monohydrochloride hydrate (Strem Chemicals, Inc., UK). Plant growth trays were rotated every day to help reduce gradient effects of light, temperature and humidity and two fully expanded young leaves from the middle of the rosette were sampled for Inductively Couple Plasma Mass Spectroscopy (ICP-MS) analysis.

#### Seed ionome

Plants were grown in 32mm 7C Jiffy-7^®^ peat pellets prepared as above. The experiment was divided into four phases: *Phase 1: Sowing* - 3 seeds of each accession were sown in pre-soaked Jiffy-7^®^ pellets, stratified at 4°C in the dark for one week. Germinated seedlings were thinned to one after one week. *Phase 2: Initial growth* - Plants were grown for two weeks in a greenhouse with supplemental lighting set at 12h day/12 h night at 26°C day/16°C night. Plants were bottom-watered once a week with 0.25 strength Hoagland solution in which Fe was supplied as 10 μM Fe-HBED. *Phase 3: Vernalisation* - Plants were placed in a cold room for eight weeks with 10 h days (50 μmol photons m^-2^s^-1^)/14 h night and temperature of 4°C day/ 2°C night. They were bottom watered twice with tap water. *Phase 4: Final growth* - Vernalised plants were returned to the greenhouse and placed randomly in 61 new trays distributed across 4 benches (**Figure S1c**). Plants were bottom watered with 0.25 strength Hoagland solution four times in the first two weeks and kept in the greenhouse for approximately two months until all seed pods were dry. Dry seeds were harvested and stored in paper envelopes. In total 6 replicates of each accession were grown and seeds harvested.

### Ionomic profiling by Inductively Couple Plasma Mass Spectroscopy (ICP-MS) analysis

#### Leaf and seed ionome

The samples were analysed using the method described by (Danku *et al*., 2013) with some modifications. For the leaf ionome, two or more leaves (2 to 5mg dry weight) were harvested per plant, washed with 18 MΩ deionized water and placed in Pyrex digestion tubes. The leaf samples were dried in the Pyrex tubes for 20h at 88°C and digested with 1 mL of concentrated nitric acid (HNO3; Baker Instra-Analyzed) in dry block heaters at 115°C for 4h. For the seed ionome between 2 to 5 mg of seeds from each plant were placed in Pyrex tubes and digested with 1.5 mL of concentrated HNO3 in dry block heaters at 115°C for 5h. After digestion, samples were diluted to 10mL with 18 MΩ Milli-Q filtered water and elemental analysis performed using an Inductively Couple Plasma Mass Spectrometer (ICP-MS) (Perkin Elmer, NexION 300D) in the standard mode and monitored for twenty-two elements (Li, B, Na, Mg, P, S, K, Ca, Cr, Mn, Fe, Co, Ni, Cu, Zn, As, Se, Rb, Sr, Mo, Cd and Pb). In order to minimise the effect of analytical drift within and between ICP-MS runs, two liquid reference solutions composed of pooled digested leaf or seed samples of the first 10 trays of the leaf and seed experiments respectively were prepared and analysed every 10 samples for all sample sets run on the ICP-MS. To compensate for the effect of variation within and between the ICP-MS runs, we used a linear model that incorporated the effect of drift within run as a covariate, and between run differences and their interaction (Holmberg and Artursson, 2004). To calculate the Limit of Detection (LOD) of the instrument per element, 40 process blanks were included in each ICP-MS sample set, (8 blanks per plant growth tray). The blanks consisted of empty tubes that went through all the sample preparation processes alongside the samples. The detection limit was calculated based on the blanks as [Mean value of a given element in the blank + (3 x Standard Deviation over all 40 blanks)] (IUPAC 2014) and samples below detection limit were set as empty. Calibration standards (with indium internal standard) were prepared from single element standards solutions.

### Data analysis

#### Weight calculation

##### Leaf ionome

For every plant growth tray the dry weight of seven reference samples were measured and used to estimate the weight and final elemental concentration of the remaining samples based on a heuristic algorithm (more info: http://www.ionomicshub.org), that uses the best quantified elements in a sample to calculate the weight and final elemental concentrations of the samples (Lahner *et al*., 2003). The approach was extended to include all four of the check-up lines Col-0, Cvi-0, Fab-2 and Ts-1, rather than just Col-0 for improved normalisation. Based on this, the dry weight of 7 samples of each of the check lines was recorded per tray. For the calculations, only the elements with a relative standard deviation of 50 or less were included. The quality of the calculated weights was verified by performing a linear regression analysis between the calculated weights of the weighed samples and their measured weight.

##### Seed ionome

ICP-MS data processing for drift correction, detection limit and samples weight calculation were performed as described above for leaf ionome data. The weight of 14 samples in each tray were measured and used for the weight calculation heuristic algorithm method. Data for all elements in both leaf and seed of all accessions analysed are available at www.ionomicshub.org at the ‘Data Exchange’.

#### Normalisation

In order to reduce the effect of external factors and to account for variation in growth conditions, we used a normalisation approach that compensates for both, variation between and within trays. First, per element, the outlier samples that were 4.5 times higher than the interquartile range across all trays were excluded from the analysis. This threshold was chosen over the more standard 1.5 times so as to not exclude accessions with an extreme phenotype from the analysis. In addition, there were idiosyncrasies observed in some trays for some elements due to, for instance, issues with the ICP-MS instrument, which caused a large number of samples in a tray to drop below the LOD. These trays were excluded based on elemental concentrations of the check-up lines i.e., if both the upper and lower quartile across all four check-up lines combined were zero. Next, to test for spatial patterns we applied a linear model with elemental concentrations as ‘response variable’, and row, column, tray and their interactions as ‘explanatory variables’ (Dragiev, Nadon and Makarenkov, 2011). Next, we modelled per element per tray with a spatial smoother using a generalized additive model (GAM) in R (Version 3.2.2; (R Core team, no date) using the package mgcv (Version 3.2.2; (Wood, 2011). We normalised for spatial patterns by subtracting the GAM predicted values for a tray from the un-normalised values of the tray and subsequently adding the overall mean predicted values. To evaluate the effectiveness of the normalisation we first tested if spatial patterns were still observable by repeating the linear model on the normalised values. Then, we analysed whether the normalisation reduced the variance among replicates of the accessions by performing a linear model that compared per element the ‘variance for all accessions of the un-normalised values’, ‘ionomic values normalised based on tray means’ and ‘spatial smoother normalised values’. After normalisation, the elemental values of the ionome were extracted per accession by calculating the Best Linear Unbiased Predictors (BLUPs) using a restricted maximum likelihood (REML) mixed model with only accessions as random effect and intercept as fixed effect (**Table 1**). The BLUPs allowed phenotypes to be predicted from genotypes which is important for fitting functional structural plant models. For the seed ionome, to compensate for variation both between and within tray variation we used the same normalisation approach described above for leaf ionomic data.

#### IonExplorer

The interactive tool was developed using R shiny, all plotting was performed using a combination of leaflet (to render the map) ggplot2 (to plot the heatmaps, histograms and pca) and plotly (to add interactive functionality and plot the radar plots). The user interface should work across most devices and browsers but has been optimised for devices with a minimum display width of 800px. The code base is publicly available at: https://bitbucket.org/ADAC_UoN/dr000081-web-service-ionome-seed-and-leaf-map/

## Acknowledgments

We thank the lab members who helped with the sowing and harvesting of the materials for this high-throughput experiment. We recognise the late Dr John Danku for performing the ICP-MS analysis for this research.

## Author Contributions

ACC, WvD, PR, AD and DES, planned and designed the research. ACC, WvD, performed experiments. ACC, WvD, PR, analysed data. PR, TG, and DES, designed the Ion Explorer and TG built the Ion Explorer. PK, AD, PS, contributed to revisions of the manuscript. ACC, WvD, PR, DES, wrote the manuscript. ACC, WvD and PR contributed equally.

## References

Alonso-Blanco, C. et al. (2016) ‘1,135 Genomes Reveal the Global Pattern of Polymorphism in Arabidopsis thaliana’, Cell, 166(2), pp. 481–491. doi: 10.1016/j.cell.2016.05.063.

Atwell, S. et al. (2010) ‘Genome-wide association study of 107 phenotypes in Arabidopsis thaliana inbred lines’, Nature. Nature Publishing Group, 465(7298), pp. 627–631. doi: 10.1038/nature08800.

Baryshnikova, A. et al. (2010) ‘Quantitative analysis of fitness and genetic interactions in yeast on a genome scale.’, Nature methods. NIH Public Access, 7(12), pp. 1017–24. doi: 10.1038/nmeth.1534.

Baxter, I. et al. (2008) ‘Variation in molybdenum content across broadly distributed populations of Arabidopsis thaliana is controlled by a mitochondrial molybdenum transporter (MOT1)’, PLoS Genet. 2008/05/06. Edited by J. Bergelson. Public Library of Science, 4(2), p. e1000004. doi: 10.1371/journal.pgen.1000004.

Baxter, I. et al. (2010) ‘A coastal cline in sodium accumulation in Arabidopsis thaliana is driven by natural variation of the sodium transporter AtHKT1;1’, PLoS Genet. 2010/11/19. Edited by G. P. Copenhaver. Public Library of Science, 6(11), p. e1001193. doi: 10.1371/journal.pgen.1001193.

Baxter, I. (2010) ‘Ionomics: The functional genomics of elements’, Briefings in Functional Genomics. Oxford University Press, 9(2), pp. 149–156. doi: 10.1093/bfgp/elp055.

Baxter, I. et al. (2012) ‘Biodiversity of Mineral Nutrient and Trace Element Accumulation in Arabidopsis thaliana’, PLoS one. Edited by P. Degryse. Public Library of Science, 7(4), pp. 1–12. doi: 10.1371/journal.pone.0035121.t001.

Baxter, I. R. et al. (2014) ‘Single-Kernel Ionomic Profiles Are Highly Heritable Indicators of Genetic and Environmental Influences on Elemental Accumulation in Maize Grain (Zea mays)’, PLoS ONE. Edited by X. Zhang. Public Library of Science, 9(1), p. e87628. doi: 10.1371/journal.pone.0087628.

Birmingham, A. et al. (2009) ‘Statistical methods for analysis of high-throughput RNA interference screens’, Nature Methods. NIH Public Access, pp. 569–575. doi: 10.1038/nmeth.1351.

Bouis, H. E. and Saltzman, A. (2017) ‘Improving nutrition through biofortification: A review of evidence from HarvestPlus, 2003 through 2016.’, Global food security. Elsevier, 12, pp. 49–58. doi: 10.1016/j.gfs.2017.01.009.

Broadley, M. R. et al. (2010) ‘An efficient procedure for normalizing ionomics data for Arabidopsis thaliana’, New Phytologist. Blackwell Publishing Ltd, 186(2), pp. 270–274. doi: 10.1111/j.1469-8137.2009.03145.x.

Buescher, E. et al. (2010) ‘Natural Genetic Variation in Selected Populations of Arabidopsis thaliana Is Associated with Ionomic Differences’, PLoS one. June 14, 2. Edited by P. K. Ingvarsson. Public Library of Science, 5(6), pp. 1–10. doi: 10.1371/journal.pone.0011081.t001.

Bus, A. et al. (2014) ‘Species- and genome-wide dissection of the shoot ionome in Brassica napus and its relationship to seedling development.’, Frontiers in plant science. Frontiers Media SA, 5, p. 485. doi: 10.3389/fpls.2014.00485.

Busoms, S. et al. (2015) ‘Salinity Is an Agent of Divergent Selection Driving Local Adaptation of Arabidopsis to Coastal Habitats’, Plant Physiology, 168(3), pp. 915–929. doi: 10.1104/pp.15.00427.

Busoms, S. et al. (2018) ‘Fluctuating selection on migrant adaptive sodium transporter alleles in coastal Arabidopsis thaliana’, Proceedings of the National Academy of Sciences of the United States of America. National Academy of Sciences, 115(52), pp. E12443–E12452. doi: 10.1073/pnas.1816964115.

Cailliatte, R. et al. (2010) ‘High-Affinity Manganese Uptake by the Metal Transporter NRAMP1 Is Essential for Arabidopsis Growth in Low Manganese Conditions’, The Plant Cell, 22(3), pp. 904–917. doi: 10.1105/tpc.109.073023.

Cao, J. et al. (2011) ‘Whole-genome sequencing of multiple Arabidopsis thaliana populations’, Nature Genetics. Nature Publishing Group, 43(10), pp. 956–963. doi: 10.1038/ng.911.

Caraus, I. et al. (2015) ‘Detecting and overcoming systematic bias in high-throughput screening technologies: a comprehensive review of practical issues and methodological solutions’, Briefings in Bioinformatics, 16(6), pp. 974–986. doi: 10.1093/bib/bbv004.

Chao, D.-Y. et al. (2011) ‘Sphingolipids in the Root Play an Important Role in Regulating the Leaf Ionome in Arabidopsis thaliana’, The Plant Cell, 23(3), pp. 1061–1081. doi: 10.1105/tpc.110.079095.

Chao, D.-Y., Chen, Y., et al. (2014) ‘Genome-wide Association Mapping Identifies a New Arsenate Reductase Enzyme Critical for Limiting Arsenic Accumulation in Plants’, PLoS Biology. Edited by J. N. Maloof. Public Library of Science, 12(12), p. e1002009. doi: 10.1371/journal.pbio.1002009.

Chao, D.-Y., Baraniecka, P., et al. (2014) ‘Variation in Sulfur and Selenium Accumulation Is Controlled by Naturally Occurring Isoforms of the Key Sulfur Assimilation Enzyme ADENOSINE 5’-PHOSPHOSULFATE REDUCTASE2 across the Arabidopsis Species Range’, Plant Physiology, 166(3), pp. 1593–1608. doi: 10.1104/pp.114.247825.

Chao, D.-Y. Y. et al. (2012) ‘Genome-wide association studies identify heavy metal ATPase3 as the primary determinant of natural variation in leaf cadmium in Arabidopsis thaliana’, PLoS Genet. 2012/09/13. Edited by K. Bomblies. Public Library of Science, 8(9), p. e1002923. doi: 10.1371/journal.pgen.1002923.

Chen, Z.-R. et al. (2018) ‘AtHMA4 Drives Natural Variation in Leaf Zn Concentration of Arabidopsis thaliana’, Frontiers in Plant Science. Frontiers, 9, p. 270. doi: 10.3389/fpls.2018.00270.

Connolly, E. L., Fett, J. P. and Guerinot, M. Lou (2002) ‘Expression of the IRT1 Metal Transporter Is Controlled by Metals at the Levels of Transcript and Protein Accumulation’, The Plant Cell Online, 14(6), pp. 1347–1357. doi: 10.1105/tpc.001263.

Danku, J. M. C. et al. (2013) ‘Large-Scale Plant Ionomics’, in Plant Mineral Nutrients. Totowa, NJ: Humana Press, pp. 255–276. doi: 10.1007/978-1-62703-152-3_17.

Dragiev, P., Nadon, R. and Makarenkov, V. (2011) ‘Systematic error detection in experimental high-throughput screening’, Bmc Bioinformatics, 12. doi: 10.1186/1471-2105-12-25.

Eide, D. J. et al. (2005) ‘Characterization of the yeast ionome: a genome-wide analysis of nutrient mineral and trace element homeostasis in Saccharomyces cerevisiae.’, Genome Biology, 6(9), p. R77. doi: 10.1186/gb-2005-6-9-r77.

Forsberg, S. K. G. et al. (2015) ‘The Multi-allelic Genetic Architecture of a Variance-Heterogeneity Locus for Molybdenum Concentration in Leaves Acts as a Source of Unexplained Additive Genetic Variance’, PLoS Genet. Edited by G. P. Copenhaver. Public Library of Science, 11(11), p. e1005648. doi: 10.1371/journal.pgen.1005648.

Gao, Y.-Q. et al. (2017) ‘A new vesicle trafficking regulator CTL1 plays a crucial role in ion homeostasis’, PLOS Biology. Edited by M. Estelle. Public Library of Science, 15(12), p. e2002978. doi: 10.1371/journal.pbio.2002978.

Guilland, C. et al. (2018) ‘Biodiversity of urban soils for sustainable cities’, Environmental Chemistry Letters. Springer Verlag, pp. 1267–1282. doi: 10.1007/s10311-018-0751-6.

Hadsell, D. L. et al. (2018) ‘In silico mapping of quantitative trait loci (QTL) regulating the milk ionome in mice identifies a milk iron locus on chromosome 1’, Mammalian Genome. doi: 10.1007/s00335-018-9762-7.

Hastie, T. and Tibshirani, R. (1986) ‘Generalized Additive Models’, Statistical Science. Institute of Mathematical Statistics, 1(3), pp. 297–310. doi: 10.1214/ss/1177013604.

Hindt, M. N. et al. (2017) ‘BRUTUS and its paralogs, BTS LIKE1 and BTS LIKE2, encode important negative regulators of the iron deficiency response in Arabidopsis thaliana’, Metallomics, 9(7), pp. 876–890. doi: 10.1039/C7MT00152E.

Holmberg, M. and Artursson, T. (2004) ‘Drift Compensation, Standards, and Calibration Methods’, in Handbook of Machine Olfaction. Weinheim, FRG: Wiley-VCH Verlag GmbH & Co. KGaA, pp. 325–346. doi: 10.1002/3527601597.ch13.

Hosmani, P. S. et al. (2013) ‘Dirigent domain-containing protein is part of the machinery required for formation of the lignin-based Casparian strip in the root.’, Proceedings of the National Academy of Sciences of the United States of America. National Academy of Sciences, 110(35), pp. 14498–503. doi: 10.1073/pnas.1308412110.

Hovick, S. M. and Whitney, K. D. (2019) ‘Propagule pressure and genetic diversity enhance colonization by a ruderal species: a multi-generation field experiment’, Ecological Monographs. Ecological Society of America, 89(3). doi: 10.1002/ecm.1368.

Huang, X.-Y. and Salt, D. E. (2016) ‘Plant Ionomics: From Elemental Profiling to Environmental Adaptation’, Molecular Plant, 9(6), pp. 787–797. doi: http://dx.doi.org/10.1016/j.molp.2016.05.003.

Huang, X.-Y. Y. et al. (2016) ‘Nuclear Localised MORE SULPHUR ACCUMULATION Epigenetically Regulates Sulphur Homeostasis in Arabidopsis thaliana’. Edited by T. Leustek. Public Library of Science, 12(9), p. e1006298. doi: 10.1371/journal.pgen.1006298.

Hussain, D. et al. (no date) ‘P-type ATPase heavy metal transporters with roles in essential zinc homeostasis in Arabidopsis’, Am Soc Plant Biol. Available at: http://www.plantcell.org/content/16/5/1327.short (Accessed: 2 October 2018).

Kamiya, T. et al. (2015) ‘The MYB36 transcription factor orchestrates Casparian strip formation.’, Proceedings of the National Academy of Sciences of the United States of America. National Academy of Sciences, 112(33), pp. 10533–8. doi: 10.1073/pnas.1507691112.

Konz, T. et al. (2017) ‘ICP-MS/MS-Based Ionomics: A Validated Methodology to Investigate the Biological Variability of the Human Ionome’, Journal of Proteome Research, 16(5), pp. 2080–2090. doi: 10.1021/acs.jproteome.7b00055.

Korshunova, Y. O. et al. (1999) ‘The IRT1 protein from Arabidopsis thaliana is a metal transporter with a broad substrate range’, Plant Molecular Biology. Kluwer Academic Publishers, 40(1), pp. 37–44. doi: 10.1023/a:1026438615520.

Kover, P. X. et al. (2009) ‘A Multiparent Advanced Generation Inter-Cross to fine-map quantitative traits in Arabidopsis thaliana’, PLoS Genetics. 2009/07/14. Edited by R. Mauricio. Public Library of Science, 5(7), p. e1000551. doi: 10.1371/journal.pgen.1000551.

Lahner, B. et al. (2003) ‘Genomic scale profiling of nutrient and trace elements in Arabidopsis thaliana’, Nature Biotechnology. 2003/09/02. Nature Publishing Group, 21(10), pp. 1215–1221. doi: 10.1038/nbt865.

Long, Q. et al. (2013) ‘Massive genomic variation and strong selection in Arabidopsis thaliana lines from Sweden’, Nat Genet. Nature Publishing Group, a division of Macmillan Publishers Limited. All Rights Reserved., 45(8), pp. 884–890. doi: 10.1038/ng.2678 http://www.nature.com/ng/journal/v45/n8/abs/ng.2678.html#supplementary-information.

Lou, M. et al. (2015) ‘Worldwide Genetic Diversity for Mineral Element Concentrations in Rice Grain’, crop science, 55. Available at: http://eprints.nottingham.ac.uk/40133/1/Pinson et al. 2015.pdf (Accessed: 24 January 2018).

Ma, S. et al. (2015) ‘Organization of the Mammalian Ionome According to Organ Origin, Lineage Specialization, and Longevity’, Cell Reports, 13(7), pp. 1319–1326. doi: 10.1016/j.celrep.2015.10.014.

Macel, M. et al. (2007) ‘Climate vs. soil factors in local adaptation of two common plant species.’, Ecology, 88(2), pp. 424–33. Available at: http://www.ncbi.nlm.nih.gov/pubmed/17479760 (Accessed: 1 February 2018).

Malinouski, M. et al. (2014) ‘Genome-wide RNAi ionomics screen reveals new genes and regulation of human trace element metabolism’, Nature Communications, 5(1), p. 3301. doi: 10.1038/ncomms4301.

Malo, N. et al. (2006) ‘Statistical practice in high-throughput screening data analysis’, NATURE BIOTECHNOLOGY VOLUME, 24(2). doi: 10.1038/nbt1186.

McDowell, S. C. et al. (2013) ‘Elemental concentrations in the seed of mutants and natural variants of Arabidopsis thaliana grown under varying soil conditions.’, PLOS one. Public Library of Science, 8(5), p. e63014. doi: 10.1371/journal.pone.0063014.

Mendoza-Cózatl, D. G. et al. (2011) ‘Long-distance transport, vacuolar sequestration, tolerance, and transcriptional responses induced by cadmium and arsenic’, Current Opinion in Plant Biology, 14(5), pp. 554–562. doi: 10.1016/j.pbi.2011.07.004.

Mills, R. F. et al. (2003) ‘Functional expression of AtHMA4, a P1B-type ATPase of the Zn/Co/Cd/Pb subclass.’, The Plant journal: for cell and molecular biology, 35(2), pp. 164–76. Available at: http://www.ncbi.nlm.nih.gov/pubmed/12848823 (Accessed: 2 October 2018).

Mills, R. F. et al. (2005) ‘The plant P _1b_-type ATPase AtHMA4 transports Zn and Cd and plays a role in detoxification of transition metals supplied at elevated levels’, FEBS Letters, 579(3), pp. 783–791. doi: 10.1016/j.febslet.2004.12.040.

Morel, M. et al. (2008) ‘AtHMA3, a P1B-ATPase Allowing Cd/Zn/Co/Pb Vacuolar Storage in Arabidopsis’, PLANT PHYSIOLOGY, 149(2), pp. 894–904. doi: 10.1104/pp.108.130294.

Morrissey, J. et al. (2009) ‘The Ferroportin Metal Efflux Proteins Function in Iron and Cobalt Homeostasis in Arabidopsis#x2019;;, The Plant Cell, 21(10), pp. 3326.LP – 3338. Available at: http://www.plantcell.org/content/21/10/3326.abstract.

Murie, C. et al. (2015) ‘Improving Detection of Rare Biological Events in High-Throughput Screens’, Journal of Biomolecular Screening. SAGE Publications Inc., 20(2), pp. 230–241. doi: 10.1177/1087057114548853.

Neugebauer, K. et al. (2018) ‘Variation in the angiosperm ionome’, Physiologia Plantarum. Blackwell Publishing Ltd, 163(3), pp. 306–322. doi: 10.1111/ppl.12700.

Nordborg, M. et al. (2005) ‘The pattern of polymorphism in Arabidopsis thaliana’, PLoS Biol. Edited by T. Mitchell-Olds. Public Library of Science, 3(7), p. e196. doi: 10.1371/journal.pbio.0030196.

Oomen, R. J. F. J. et al. (2009) ‘Functional characterization of NRAMP3 and NRAMP4 from the metal hyperaccumulator Thlaspi caerulescens’, New Phytologist. Blackwell Publishing Ltd, 181(3), pp. 637–650. doi: 10.1111/j.1469-8137.2008.02694.x.

Pauli, D. et al. (2018) ‘Multivariate Analysis of the Cotton Seed Ionome Reveals a Shared Genetic Architecture.’, G3 (Bethesda, Md.). G3: Genes, Genomes, Genetics, 8(4), pp. 1147–1160. doi: 10.1534/g3.117.300479.

Poormohammad Kiani, S. et al. (2012) ‘Allelic Heterogeneity and Trade-Off Shape Natural Variation for Response to Soil Micronutrient’, PLoS Genetics. Edited by R. Mauricio. Public Library of Science, 8(7), p. e1002814. doi: 10.1371/journal.pgen.1002814.

R Core team (no date) R: a language and environment for statistical computing, Vienna, Austria, 2018.

Ricachenevsky, F. K. et al. (2018) ‘You Shall Not Pass: Root Vacuoles as a Symplastic Checkpoint for Metal Translocation to Shoots and Possible Application to Grain Nutritional Quality.’, Frontiers in plant science, 9, p. 412. doi: 10.3389/fpls.2018.00412.

Rus, A. et al. (2006) ‘Natural Variants of AtHKT1 Enhance Na+ Accumulation in Two Wild Populations of Arabidopsis’, PLoS Genetics. Public Library of Science, 2(12), p. e210. doi: 10.1371/journal.pgen.0020210.

Salt, D. E., Baxter, I. and Lahner, B. (2008) ‘Ionomics and the Study of the Plant Ionome’, Annual Review of Plant Biology. Annual Reviews, 59(1), pp. 709–733. doi: 10.1146/annurev.arplant.59.032607.092942.

Takou, M. et al. (2019) ‘Linking genes with ecological strategies in Arabidopsis thaliana’, Journal of Experimental Botany. Oxford Academic, 70(4), pp. 1141–1151. doi: 10.1093/JXB/ERY447.

Thomas, C. L. et al. (2016) ‘Root morphology and seed and leaf ionomic traits in a Brassica napus L. diversity panel show wide phenotypic variation and are characteristic of crop habit’, BMC Plant Biology. BioMed Central, 16(1), p. 214. doi: 10.1186/s12870-016-0902-5.

Tian, H. et al. (2010) ‘Arabidopsis NPCC6/NaKR1 is a phloem mobile metal binding protein necessary for phloem function and root meristem maintenance.’, The Plant cell. American Society of Plant Biologists, 22(12), pp. 3963–79. doi: 10.1105/tpc.110.080010.

Verret, F. et al. (2004) ‘Overexpression of AtHMA4 enhances root-to-shoot translocation of zinc and cadmium and plant metal tolerance’, FEBS Letters, 576(3), pp. 306–312. doi: 10.1016/j.febslet.2004.09.023.

Visscher, P. M., Hill, W. G. and Wray, N. R. (2008) ‘Heritability in the genomics era — concepts and misconceptions’, Nature Reviews Genetics, 9(4), pp. 255–266. doi: 10.1038/nrg2322.

Vreugdenhil, D. et al. (2004) ‘Natural variation and QTL analysis for cationic mineral content in seeds of Arabidopsis thaliana’, Plant, Cell and Environment, 27(7), pp. 828–839. doi: 10.1111/j.1365-3040.2004.01189.x.

Watanabe, T. et al. (2007) ‘Evolutionary control of leaf element composition in plants’, New Phytologist, 174(3), pp. 516–523. doi: 10.1111/j.1469-8137.2007.02078.x.

Waters, B. M. and Sankaran, R. P. (2011) ‘Moving micronutrients from the soil to the seeds: Genes and physiological processes from a biofortification perspective’, Plant Science, 180(4), pp. 562–574. doi: 10.1016/j.plantsci.2010.12.003.

Weigel, D. and Mott, R. (2009) ‘The 1001 Genomes Project for Arabidopsis thaliana’, Genome Biology, 10(5), pp. 1–5. doi: 10.1186/gb-2009-10-5-107.

White, P. J. (2001) ‘The pathways of calcium movement to the xylem’, Journal of Experimental Botany. Oxford University Press, 52(358), pp. 891–899. doi: 10.1093/jexbot/52.358.891.

Wiles, A. M. et al. (2008) ‘An Analysis of Normalization Methods for *Drosophila* RNAi Genomic Screens and Development of a Robust Validation Scheme’, Journal of Biomolecular Screening, 13(8), pp. 777–784. doi: 10.1177/1087057108323125.

Wintz, H. et al. (2003) ‘Expression Profiles of *Arabidopsis thaliana* in Mineral Deficiencies Reveal Novel Transporters Involved in Metal Homeostasis’, Journal of Biological Chemistry, 278(48), pp. 47644–47653. doi: 10.1074/jbc.M309338200.

Wood, S. N. (2011) ‘Fast stable restricted maximum likelihood and marginal likelihood estimation of semiparametric generalized linear models’, Journal of the Royal Statistical Society. Series B: Statistical Methodology, 73(1), pp. 3–36. doi: 10.1111/j.1467-9868.2010.00749.x.

Yang, M. et al. (2018) ‘Genome-wide association studies reveal the genetic basis of ionomic variation in rice’, Plant Cell. American Society of Plant Biologists, 30(11), pp. 2720–2740. doi: 10.1105/tpc.18.00375.

Yoneyama, T., Ishikawa, S. and Fujimaki, S. (2015) ‘Route and Regulation of Zinc, Cadmium, and Iron Transport in Rice Plants (Oryza sativa L.) during Vegetative Growth and Grain Filling: Metal Transporters, Metal Speciation, Grain Cd Reduction and Zn and Fe Biofortification’, International Journal of Molecular Sciences, 16(8), pp. 19111–19129. doi: 10.3390/ijms160819111.

Yu, D. et al. (2011) ‘Noise reduction in genome-wide perturbation screens using linear mixed-effect models’, Bioinformatics, 27(16), pp. 2173–2180. doi: 10.1093/bioinformatics/btr359.

Yu, D. et al. (2012) ‘High-resolution genome-wide scan of genes, gene-networks and cellular systems impacting the yeast ionome’, BMC Genomics. BMC Genomics, 13(1), p. 623. doi: 10.1186/1471-2164-13-623.

Ziegler, G. et al. (2018) ‘Genomewide association study of ionomic traits on diverse soybean populations from germplasm collections’, Plant Direct, 2(1). doi: 10.1002/PLD3.33.

