## Supplemental data for "Full species-wide leaf and seed ionomic diversity of *Arabidopsis thaliana*"

[1]Institute of Biological and Environmental Sciences, University of Aberdeen, Cruickshank Building, Aberdeen, AB24 3UU, United Kingdom; [2]Future Food Beacon of Excellence and School of Biosciences, University of Nottingham, Sutton Bonington Campus, Loughborough, Leicestershire, LE12 5RD, United Kingdom; [3]Digital Research Service and Advanced Data Analysis Centre, University of Nottingham; Sutton Bonington Campus, Loughborough, Leicestershire, LE12 5RD, United Kingdom; [4]Gregor Mendel Institute of Molecular Plant Biology, Vienna, Austria.

*Corresponding author: Professor David E Salt

[#]Authors contributed equally

### Supplementary Figures

**Figure S1** a) Radar plot representing elemental leaf ionome profile of the *Arabidopsis thaliana* accessions Col-0, Ts-1, Cvi-0 and Fab-2 that were used as to normalise the experimental data. Axis display Z-scores calculated per element. b) Distribution of normalisation lines over trays in the leaf experiments. C) Randomisation map of normalisation lines used for seed ionome experiment (c). Blue, orange, yellow and red plants indicate the different normalisation lines.

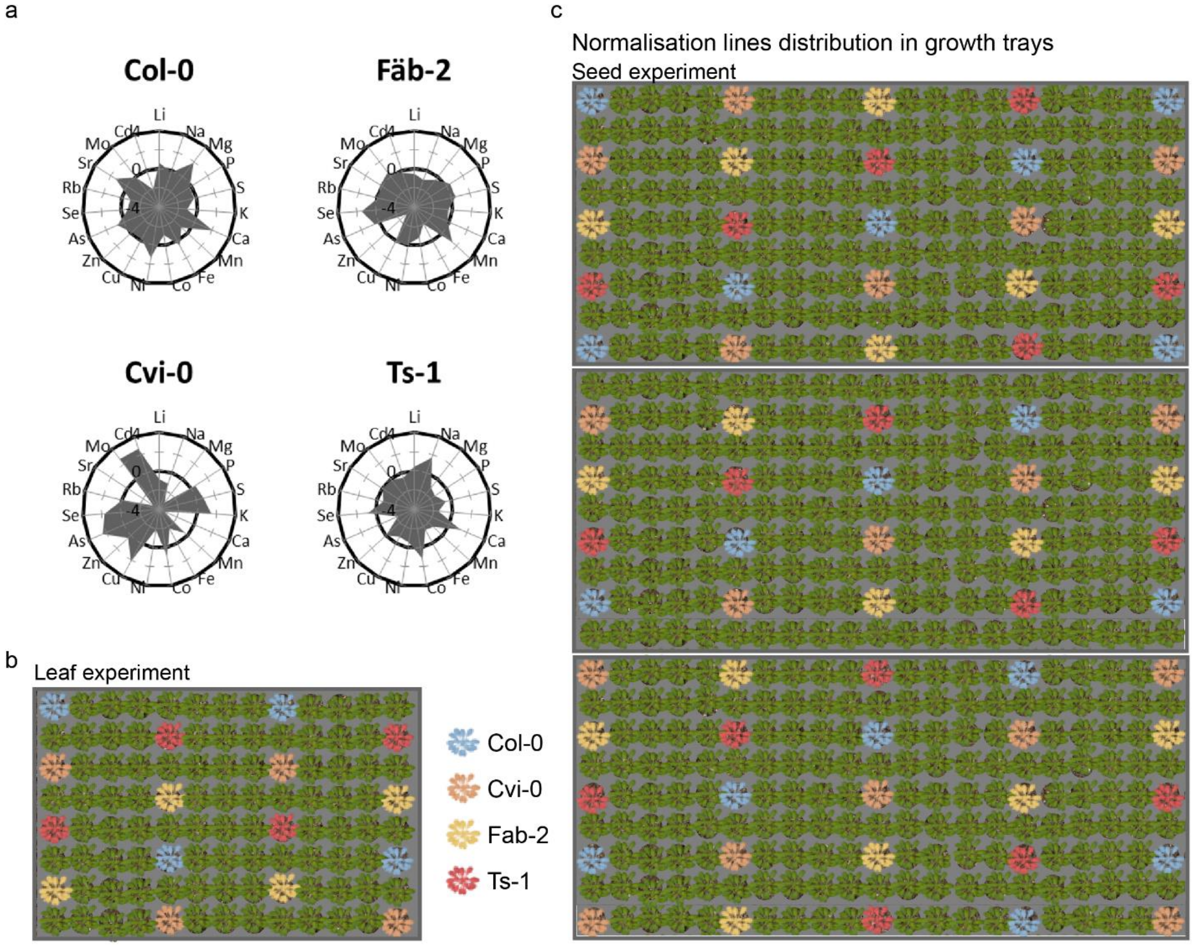

**Figure S2** Distribution of un-normalised Se concentrations over all samples per tray (a) and check-up lines (b) per tray. Distribution of Se concentration of all samples (c) and check-up lines (d) per trays after normalisation.

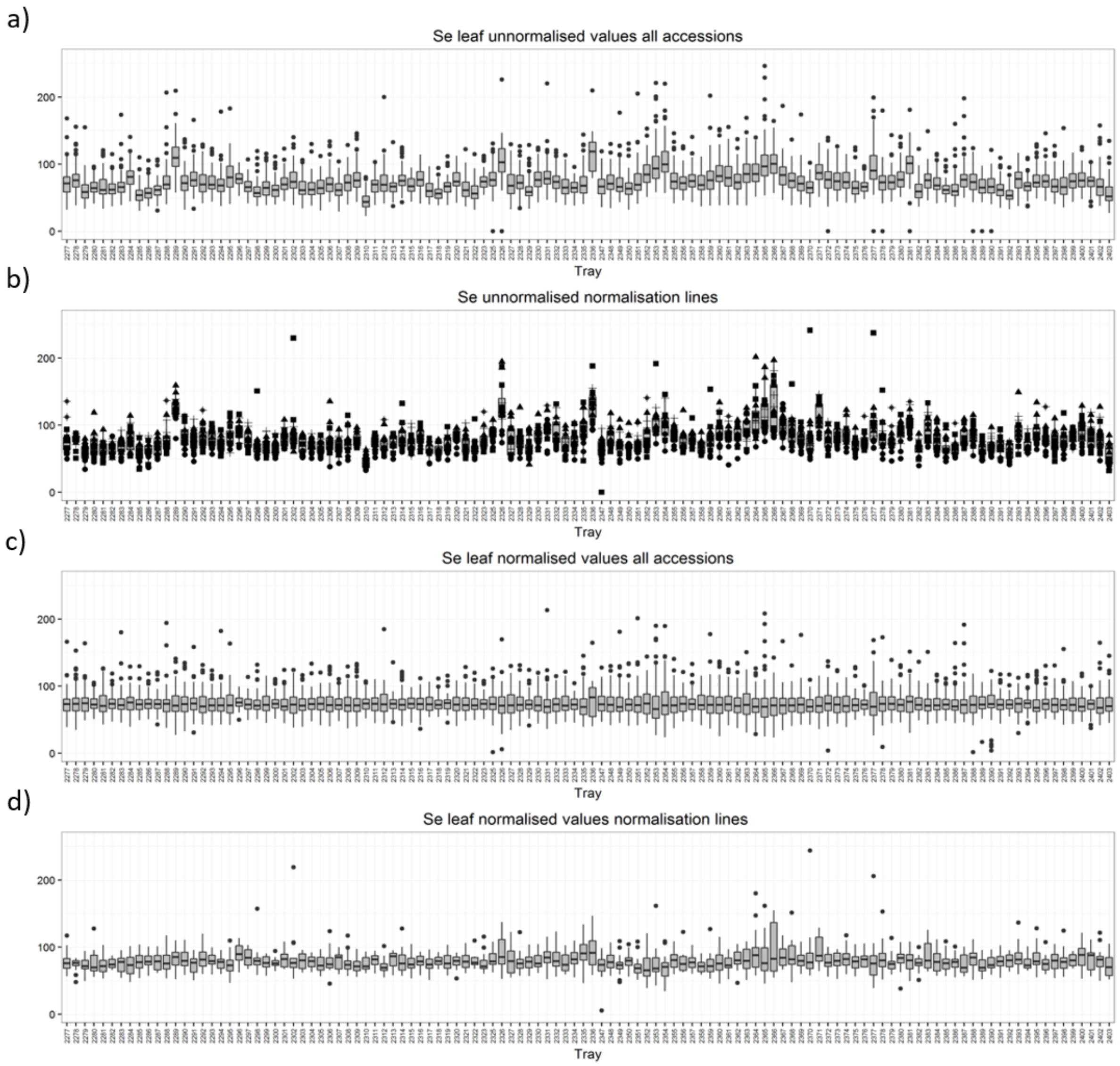

**Figure S3** (a) Un-normalised Se concentrations of individual samples across trays, (b) Predicted values of Se concentration across trays based on a GAM, (c) Normalised Se concentration of individual samples across trays based on the predicted values obtained from a GAM.

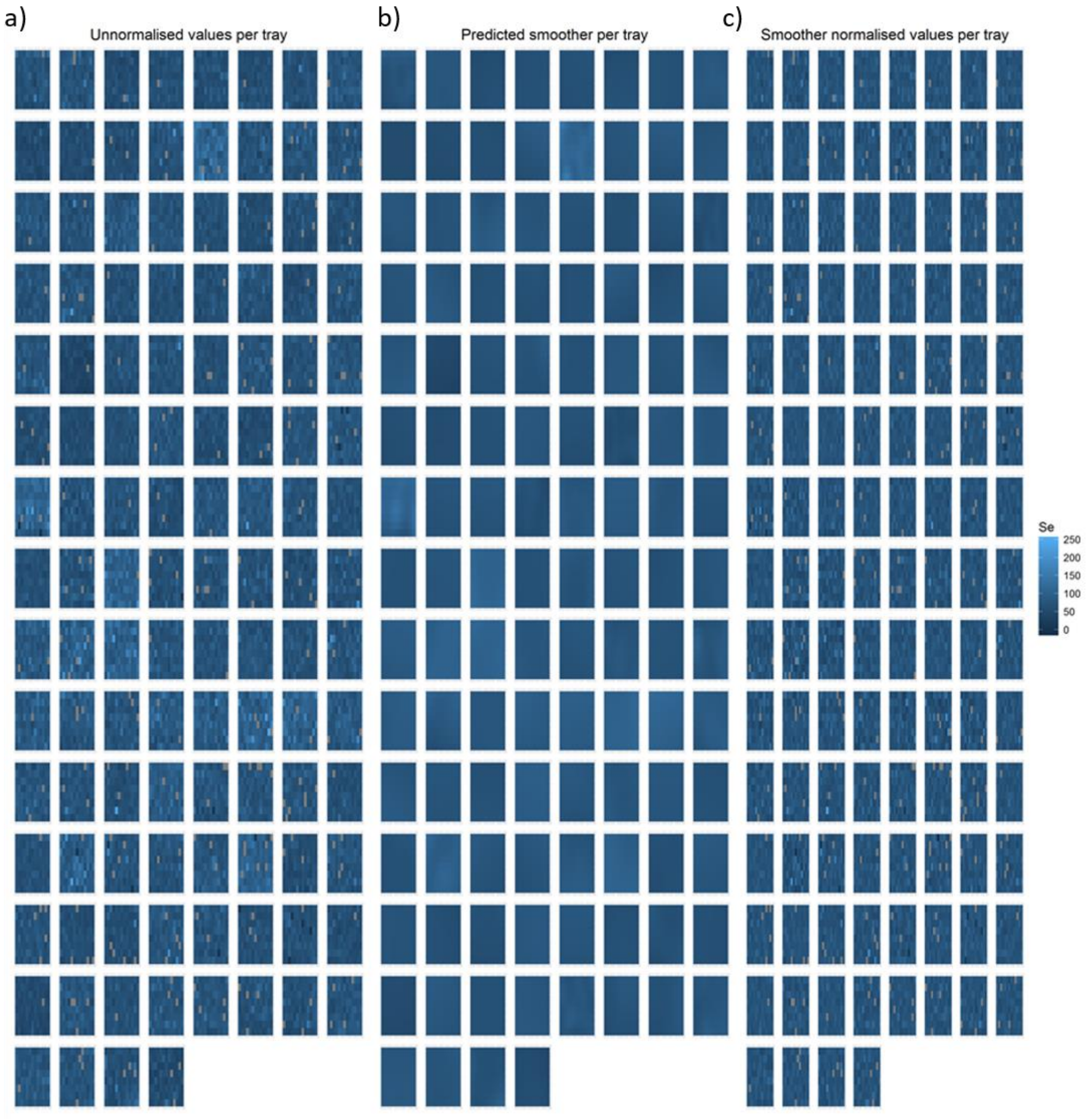

**Figure S4** Biplot of the Principle Component Analysis of the leaf (a) and seed ionome (b) of the studied *A. thaliana* accessions grouped based on geographical parameters in Alonso Blanco *et al*., 2016.

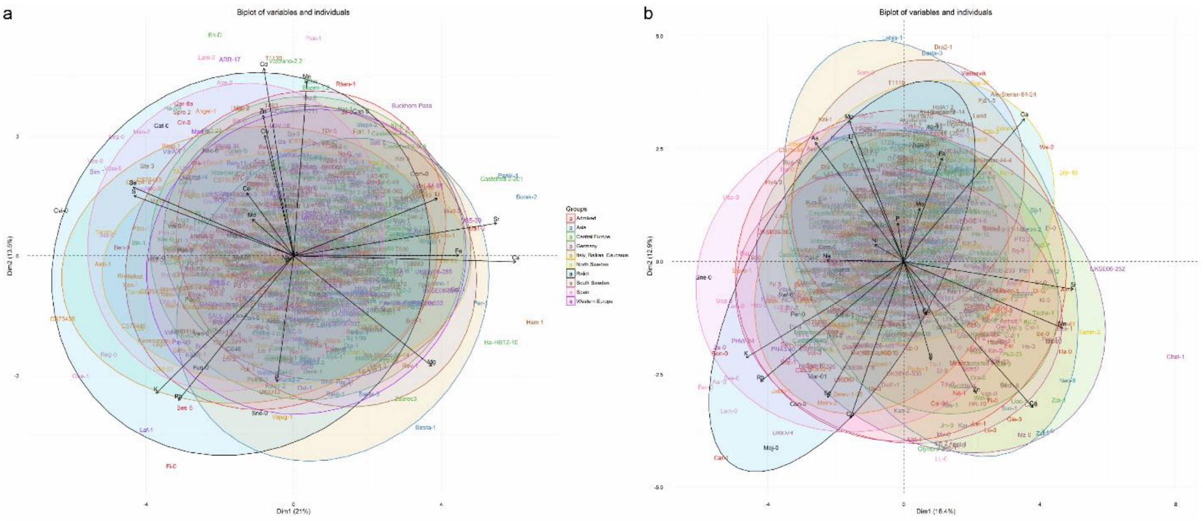


**Figure S5** Heatmap that shows the hierarchical clustering of analysed accessions based on their leaf (a) and seed (b) ionome. The top row represents the regional groups based on genetic similarities and geographic distance (Alonso-Blanco *et al*., 2016).

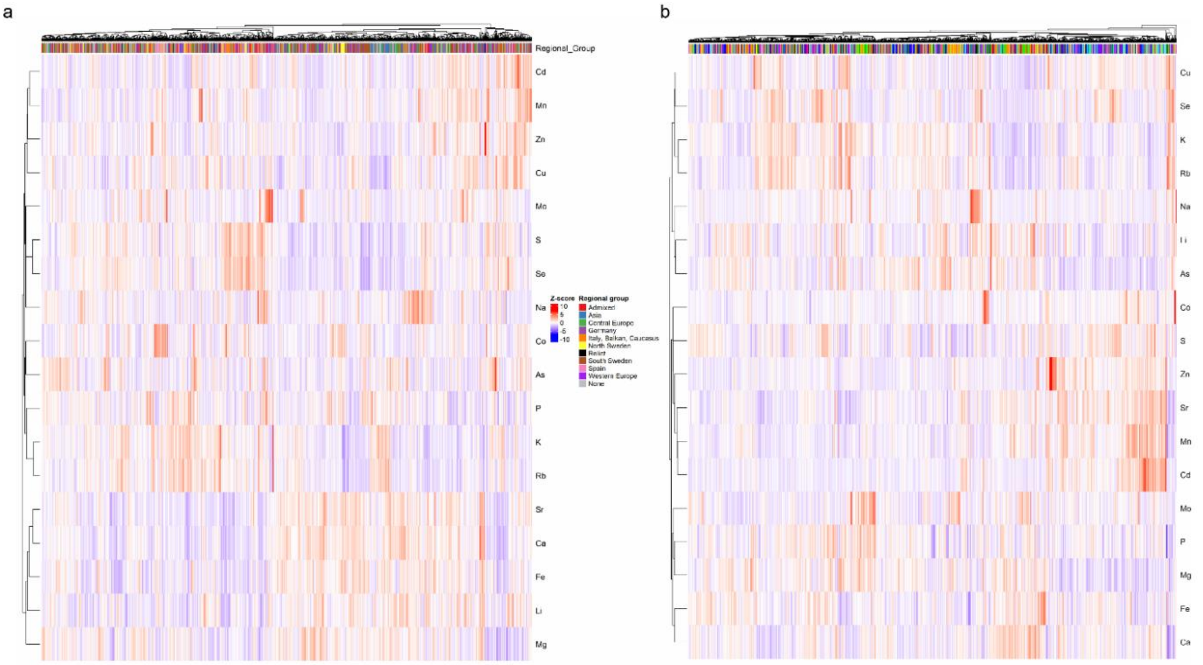

**Figure S6** Heatmap showing the hierarchical clustering of the accessions studied based on ten top and bottom extreme accessions for the leaf (a) and seed (b) ionome. The top row represents the regional groups based on genetic similarities and geographic distances (Alonso-Blanco *et al.*, 2016).

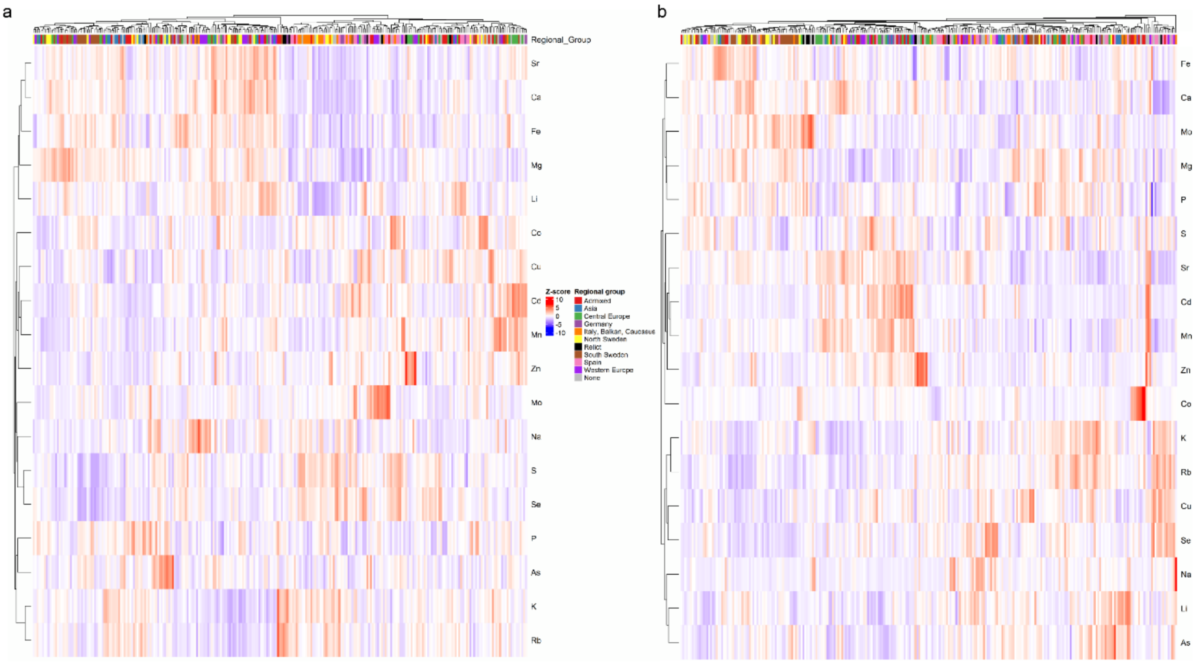

### Supplementary tables

**Table S1. Output of the linear model on row, column, tray and their interaction on the leaf ionome of the accessions studied.** R*C: Row*Column, R*T: Row*Tray, C*T: Column*Tray, R*C*T: Row*Column*Tray. Significance levels in black are derived from a linear model on the unnormalized ionomic values, significance levels in red are derived from the same linear model on the normalized ionomic values. Significance levels: ***:p<0.001, **:p<0.01, *:p<0.05.

| Element | Row | Column | Tray | R*C | R*T | C*T | R*C*T |
|---|---|---|---|---|---|---|---|
| **Li** | *** | | *** | | *** | *** | *** |
| **Na** | ** | *** | *** | *** | * | *** | * |
| **Mg** | | | *** | *** | *** | *** | |
| **P** | | ** | *** | * | | *** | |
| **S** | *** | *** | *** | * | *** | *** | *** |
| **K** | | *** | *** | | *** | *** | * |
| **Ca** | | *** | *** | *** | *** | *** | |
| **Mn** | | *** | *** | | | *** | |
| **Fe** | *** | *** | *** | | *** | *** | *** |
| **Co** | | ** | *** | *** | *** | ** | |
| **Cu** | | | *** | | | *** | ** |
| **Zn** | | *** | *** | *** | | *** | *** |
| **As** | | ** | *** | * | *** | *** | |
| **Se** | * | *** | *** | *** | | *** | |
| **Rb** | ** | | *** | | | *** | |
| **Sr** | | *** | *** | *** | *** | *** | *** |
| **Mo** | | * | *** | | | | |
| **Cd** | | | *** | | | ** | |

**Table S2 Output of the linear model on row, column, tray and their interaction on the seed ionome of the accessions studied. R*C: Row*Column, R*T: Row*Tray, C*T:** Column*Tray, R*C*T: Row*Column*Tray. Significance levels in black are derived from a linear model on the un-normalised ionomic values, significance levels in red are derived from the same linear model on the normalised ionomic values. Significance levels: ***:p<0.001, **:p<0.01, *:p<0.05.

| **Element** | **Row** | **Column** | **Tray** | **R*C** | **R*T** | **C*T** | **R*C*T** |
|---|---|---|---|---|---|---|---|
| **Li** | *** | ** | *** | | *** | *** | * |
| **Na** | | | *** | | ** | *** | |
| **Mg** | | *** | *** | | ** | *** | |
| **P** | | *** | *** | | *** | *** | ** |
| **S** | *** | *** | *** | *** | *** | *** | ** |
| **K** | ** | * | *** | | *** | *** | |
| **Ca** | | | *** | | *** | *** | |
| **Mn** | *** | *** | *** | | *** | *** | |
| **Fe** | | *** | *** | * | *** | *** | |
| **Co** | | * | *** | | | *** | |
| **Ni** | | | *** | | | ** | |
| **Cu** | | * | *** | | *** | *** | |
| **Zn** | | *** | *** | | | *** | |
| **As** | *** | *** | *** | | *** | *** | |
| **Se** | | | *** | | *** | *** | ** |
| **Rb** | | * | *** | | *** | *** | * |
| **Sr** | *** | | *** | | *** | *** | * |
| **Mo** | *** | | *** | | * | * | |
| **Cd** | | *** | *** | | *** | *** | |

**Table S3 Overview of the ten accessions with highest and lowest concentration levels of each element measured in leaves across the studied *A. thaliana* accessions.** Accessions that occur multiple times are highlighted in bold; Check-up line and MAGIC line parents that appear are in red

| Element | Bottom 10 accessions | Top 10 accessions |
|---|---|---|
| **Li** | **UllA 1**, **CS75423**, Tamm-27, Tamm-2, CS75432, **Lago-1**, **Reg-0**, Apost-1, Xan-1, CS75419 | **Ba-1**, **Ezc-2**, **Cnt-1**, Gy-0, Var2-1, Pna-17, **Castelfed-2-201**, Vpa-1, UKSE06-118, Sac-0 |
| **Na** | PHW-34, **Ini-0**, Aul-0, Fell3-7, TFÄ 06, LCL-16, Kolyv-3, UKSW06-207, **Ha-SB**, Brest-1 | CS75444, BEZ-9, UKNW06-003, T990, TRE-1, Duk, **UKID13**, Ste 4, Mitterberg-2-185, **MNF-Pot-21** |
| **Mg** | Ele-0, **UllA 1**, Rel-0, **VarA 1**, Ala-0, Cvi-0, Lam-0, Spro 2, Coa-0, **Ha-SB** | Liarum, TV-38, TDr-18, Lan 1, St-0, DraIV 6-13, Ham 1, Kelsterbach-4, Bch-1, Ciste-2 |
| **P** | Buckhorn Pass, **Bisig-1**, Uk-3, ZdrI 1-23, Star-8, **VarA 1**, Vav-0, Ler-0, Stu1-1, Grivo-1 | Pro-0, Vajug-1, **UKID13**, Karag-2, CYR, TAA 17, TAA 04, Moa-0, UKID79, Sne-0 |
| **S** | **Ini-0**, **T1160**, T960, Rev-1, Zdarec3, T880, T840, T800, Yst 1, TDr-18 | CS75436, Fri 1, Tnz-1, CS75437, Istisu-1, Vas-0, Stilo-1, CS75416, CS75407, Hod |
| **K** | Bar 1, Eden-1, Stilo-1, Nas 2, Kia 1, **Castelfed-2-201**, TOM 01, Gud-3, T†D 05, ISS-20 | Fi-0, Me-0, Gua-1, Wa-1, Vdt-0, Fun-0, Wl-0, Ovi-1, Tha-1, Toufl-1 |
| **Ca** | **UllA 1**, Cvi-0, Vdm-0, Aiell-1, CS75436, Ala-0, Rld-1, **Reg-0**, TGR 01, **Sim 1** | Borsk-2, Ba1-2, **Castelfed-2-201**, Ha-HBT2-10, ISS-20, Adal 1, TOM 03, Mdd-0, LP3413.41, Smolj-1 |
| **Mn** | UKID116, Ws-2, Krot-0, PLY-20, Sapporo-0, Bes-5, Omn-5, **Reg-0**, Rev-1, CS75436 | Fr-2, Pva-1, ARR-17, T1000, En-D, Boa-0, Pa-1, Wil-1, Bozen-1.2, Nc-1 |
| **Fe** | **Sim 1**, Cvi-0, Slavi-2, Pra-6, **Timpo-1**, **Cho-0**, Had-1b, RAD-21, Tamm-27, **Pdl-0** | Ham-7-233, UduI 3-36, Ham 1, Fja 1-2, OMo2-3, T670, Ber, **Schip-1**, KZ10, Pva-1 |
| **Co** | Pie-0, DraIV 5-12, Mitterberg-2-185, Ste-0, **Pdl-0**, Krot-0, Altai-5, Aitba-2, **UllA 1**, Wank-2 | Stilo-1, Lerik1-3, Usa-0, UKNW06-281, Filet-1, Pue-0, Pva-1, Tam-0, TGR 02, TOM 01 |
| **Cu** | Lu-1, Basta-2, VED-10, Aln-30, Adc-5, CS75426, CS75431, Koz-2, Marti-1, Uk-1 | Angso-80-432, Lam-0, Wa-1, Ele-0, Gra-0, Vastervik, ARR-17, RRS-7, **Lag2.2**, Ey15-2 |
| **Zn** | Fab-2, CS75469, Be-0, TAL 03, GEN-8, Fun-0, NOZ-6, Fab-4, TAD 05, Ale-Stenar-56-14 | UKNW06-403, Cat-0, **MNF-Pot-21**, T1130, Tamm-2, Borsk-2, BI-4, Pu2-7, **Timpo-1**, Stw-0 |
| **As** | Tri-0, **Reg-0**, Vdt-0, UKID74, Marti-1, Omn-5, UKSE06-325, Knox-10, T570, Ang-0 | UKSE06-252, Sij-4, TBO 01, Uk-1, Bil-5, Cvi-0, PYL-6, Hau-0, Hov3-2, Cimin-1 |
| **Se** | T840, Rev-1, **T1160**, T780, T670, T880, App1-16, T690, T960, T800 | CS75436, Ace-0, Tnz-1, **Ba-1**, Vas-0, Hod, Men-2, CS75437, **Bisig-1**, CS28208 |

| | | |
|---|---|---|
| **Rb** | T990, Nas 2, Ull3-4, Eden-2, Stilo-1, **Castelfed-2-201**, Fri 2, T460, Castelfed-2-200, RUM-20 | Fi-0, **UllA 1**, Ven-0, T1090, Gua-1, Fun-0, MNF-Pin-39, Eds-1, Krepo-1, Se-0 |
| **Sr** | Cvi-0, **UllA 1**, Aiell-1, **Cho-0**, **Lago-1**, TGR 01, Pigna-1, **Sim 1**, Boa-0, Leg-0 | Panik-1, TOM 03, VED-10, Ha-HBT2-10, Teiu-1, **Castelfed-2-201**, Krazo-1, Ham 1, Castelfed-4-214, Jim-1 |
| **Mo** | Kolyv-6, CS75474, Rod-17-319, Bela-2, Lebja-2, Basta-1, Masl-1, CS75469, Krazo-1, Gr-1 | Bea-0, Cor-0, CS75653, Cdo-0, Vim-0, Ala-0, UKSW06-226, UKSW06-333, ESP-1-11, **Lag2.2** |
| **Cd** | TU-PK-7, Mitterberg-3-187, Bes-5, UKSE06-533, Aln-30, Lerik1-3, Eden-7, **UKID13**, CS75443, Mrk-0 | Vezzano-2.2, T1130, Bozen-1.2, Spro 2, **Ha-SB**, Gu-0, HSm, En-D, **Schip-1**, Bela-3 |

**Table S4 Overview of the ten accessions with highest and lowest concentration levels of each element measured in seeds across analysed *A. thaliana* accessions.** Accessions that occur multiple times are highlighted in bold; Check-up line and MAGIC line parents that appear are in red

| Element | Bottom 10 accessions | Top 10 accessions |
|---|---|---|
| **Li** | **Dolna-1-10**, Wl-0, Jm-0, Tscha-1, Ale1-2, Bsch-0, Ak-1, RAD-21, Dolna-1-40, Knjas-1 | Som-0, Sparta-1, CS75452, Anz-0, Lebja-1, Basta-3, FlyA 3, Eden-2, Pva-1, Basen-1 |
| **Na** | Ruma-1-27, Dra-0, TÅD 03, Aledal-6-49, Hi-0, Jm-0, Bran-1, CS75439, Bsch-0, Nyl 13 | CS75444, Rü-N2, UKNW06-281, Hec-0, Vas-0, Slc-3, Bil-5, Lro-0, Vis-0, Spr1-2 |
| **Mg** | Aln-30, Ovi-1, Moj-0, Mz-0, Urd-1, CHA-41, Bg-2, Anholt-1, UKID13, Ha-HBT1-2 | Staro-1, Aledal-6-49, Pal-0, T720, T1080, Iso-4, Kni-1, Västervik, T880, Castelfed-1-197 |
| **P** | Bar-1, Lam-0, Pee-0, Pa-1, Urd-1, Hue-3, Slavi-2, Ha-HBT3-11, Ler-1, **Dolna-1-10** | FlyA 3, Nac-0, Iso-4, Dja 1, TAA 04, App1-16, Sen-0, Qar-8a, EdJ 2, **Pdl-0** |
| **S** | T960, Hue-3, Bar-1, DraIV 3-7, MAR2-3, Bor-1, Ren-1, DraII-6, T850, Ste 4 | Fi-0, Uod-1, Bik-1, Pa-1, Eds-9, Had-1b, Alst-1, Mz-0, Deh-1, VårA 1 |
| **K** | CS75447, Ak-1, UKSE06-252, Lip-0, TDr-8, Sij-2, **Castelfed-2-201**, Gra-0, DraIV 3-7, UKNW06-233 | Sne-0, Aln-30, Bela-3, Lum-0, Lch-0, Voz-0, Moz-0, Car-1, Aru-0, Esn-2 |
| **Ca** | Esn-2, TV-7, Aln-30, Lam-0, **Aul-0**, Scm-0, Car-1, Vår2-1, Ses-0, UKID74 | Dör-10, Tnz-1, Chat-1, Lund, T840, Ven-0, Eden-9, Cerv-1, **Västervik**, Nyl-2 |
| **Mn** | Lam-0, Vpa-1, FlyA 3, **Aul-0**, Pal-0, Lis-2, Urd-1, Bela-3, Sen-0, Bis-0 | Lip-0, Kz-9, Pa-1, **Cnt-1**, Chat-1, Pro-0, Mz-0, Kondara, Lz-0, Pna-17 |
| **Fe** | **Cnt-1**, Est-1, Ven-0, Urd-1, Sim 1, Staro-1, Ste 2, Gra-0, UKNW06-281, Udu-12 | Bar-1, Sei-0, Stepn-1, T470, Krazo-2, EkS 2, Ale-Stenar-44-4, Eds-9, T840, Lab-7 |

| | | |
|---|---|---|
| **Co** | Mandr-1, Sfb-6, La-0, Bela-1, Amu-0, Cdo-0, Wt-5, LP3413.41, **Pdl-0**, Ko-2 | UKID67, UKSE06-325, **CS75423**, Mah-6, Fr-2, Ivano-1, Rev-0, ÖMö1-7, Mon-5, Grön-5 |
| **Ni** | Np-0, Tos-82-387, TÅD 03, UKNW06-481, Vaz-0, Alst-1, UKNW06-354, Star-8, Bå1-2, Xan-1 | T980, Gn2-3, T1080, T550, BEZ-9, Bes-5, Rü4-16, Tha-1, Pil-0, Bil-5 |
| **Cu** | Ven-0, Fjä1-5, Basta-2, Mdc-0, T850, Sei-0, TÅD 01, Som-0, T460, Altai-5 | Brest-1, Hov1-10, Aln-30, Sp-0, Scm-0, Moj-0, Lch-0, LL-0, Lam-0, Pro-0 |
| **Zn** | Kolyv-5, Usa-0, T780, Hue-3, Bis-0, Balan-1, TDr-17, Vae-2, T800, Böt 1 | Bg-5, Pa-1, Olympia-2, UduI 1-11, Tsu-0, Röd-17-319, Aa-0, Leska-1-78, Chat-1, Ven-0 |
| **As** | Vpa-1, Pi-0, Men-2, Ga-0, Del-10, Ms-0, Tamm-2, Mer-6, Pro-0, Kyr-1 | Pva-1, Som-0, Usa-0, Coy-0, Fri 3, **Ezc-2**, Sparta-1, Ty-0, Sne-0, Eden-2 |
| **Se** | TDr-17, TÅD 04, Ale-Stenar-44-4, Gul1-2, T1080, Spr1-6, Sku-30, Ale-Stenar-56-14, Tomegap-2, Dra2-1 | Cimin-1, Aln-30, Vid-1, Bae-0, Bea-0, Coa-0, Qar-8a, Sca-0, Ses-0, Zu-0 |
| **Rb** | UKSE06-252, T750, Dra2-1, Spr1-6, Lillö-1, Iso-4, Ör-1, TDr-8, UKNW06-233, Knjas-1 | **Aul-0**, Elp-0, Corig-1, Lum-0, Car-1, Kardz-2, Draha2, Zdarec3, UKSE06-432, Sne-0 |
| **Sr** | Ren-6, Bela-3, Vash-1, Aln-30, Bar-1, CS75465, VårA 1, Val-0, Ses-0, Staro-1 | Dör-10, Chat-1, Di-G, Gie-0, Ws-2, Lip-0, Kl-5, Boot-1, Pa-1, UKSE06-500 |
| **Mo** | K-oze-3, Ler-1, Sever-1, Lebja-1, HE1, Stara-1, Nosov-1, CS75469, Rubeznhoe-1, Krazo-2 | UKNW06-212, Pun-0, Iso-4, Vis-0, Fäb-4, Ale-Stenar-64-24, Cal-0, Zar-0, Castelfed-3-205, Bela-3 |
| **Cd** | Pog-0, Spro 2, CS75431, App1-16, Tur 4, CS75475, Knjas-1, Liri-1, **CS75423**, Rü4-16 | Lip-0, Zal-1, Ha-0, Pa-1, Sus-1, Gr-1, Zdr-1, Ka-0, Le-0, Kyr-1 |

### References


Alonso-Blanco, C. *et al.* (2016) '1,135 Genomes Reveal the Global Pattern of Polymorphism in Arabidopsis thaliana', *Cell*, 166(2), pp. 481–491. doi: 10.1016/j.cell.2016.05.063.
